# Fast temporal dynamics and causal relevance of face processing in the human temporal cortex

**DOI:** 10.1101/416214

**Authors:** Jessica Schrouff, Omri Raccah, Sori Baek, Vinitha Rangarajan, Sina Salehi, Janaina Mourão-Miranda, Zeinab Helili, Amy L. Daitch, Josef Parvizi

**Affiliations:** Laboratory of Behavioral and Cognitive Neuroscience, Department of Neurology and Neurological Sciences, Stanford University, California, USA; Computer Science Department, University College London, Gower street, London WC1E6BT, UK

**Author notes:** **Corresponding author:** Josef Parvizi MD PhD, 300 Pasteur drive, Palo Alto, CA 94305.

**Keywords:** Electrocorticography, Modular processing, Distributed Processing, Local Field Potentials, Spatiotemporal Dynamics, Machine Learning Classification

## Abstract

Recordings with a large number of intracranial electrodes in eight neurosurgical subjects offered a unique opportunity to examine the fast temporal dynamics of face processing simultaneously across a relatively large extent of the human temporal cortex (TC). Measuring the power of slow oscillatory bands of activity (θ, α, β, and γ) as well as High-Frequency Broadband (HFB, 70-177 Hz) signal, we found that the HFB showed the strongest univariate and multivariate changes in response to face compared to non-face stimuli. Using the HFB signal as a surrogate marker for local cortical engagement, we identified recording sites with selective responses to faces that were anatomically consistent across subjects and responded with graded strength to human, mammal, bird, and marine animal faces. Importantly, the most face selective sites were located more posteriorly and responded earlier than those with less selective responses to faces. Using machine learning based methods, we demonstrated that a sparse model focusing on information from the human face selective sites performed as well as, or better than, anatomically distributed models of face processing when discriminating faces from non-faces stimuli. Lastly, we identified the posterior fusiform (pFUS) site as causally the most relevant node for inducing distortion of face perception by direct electrical stimulation. Our findings support the notion of face information being processed first in the most selective sites - that are anatomically discrete and localizable within individual brains and anatomically consistent across subjects – which is then distributed in time to less selective anterior temporal sites within a time window that is too fast to be detected by current neuroimaging methods. The new information about the fast spatio-temporal dynamics of face processing across multiple sites of the human brain provides a new common ground for unifying the seemingly contradictory modular and distributed models of face processing in the human brain.

## INTRODUCTION

Studies using lesion methods(1, 2), functional imaging tools(3-6) or scalp encephalography (EEG) (7) and magnetoencephalography (MEG)(8, 9) have offered invaluable causal, spatial, and temporal information about the neural mechanisms of face processing in the human brain. Work in non-human primates (10-15) has also provided important novel insights. However, despite great progress, the long lasting controversy between modular versus distributed models of face processing has persisted in the literature (16). Some studies have revealed face-selective responses only in anatomically consistent regions of the temporal cortex (TC) (5), and other observations have shown that the pattern of responses to face stimuli can be discerned from sampled data from non–selective regions of the TC (17) suggesting that face information is anatomically distributed. Both theories have unfortunately relied on information with limited temporal resolution averaged over multiple seconds or from methods using regions of interest and averaging across subjects, or direct recordings from a single or a pair of recording sites. Thus, *the fast temporal dynamics of face processing across a large extent of the cerebral cortex within individual brains remains poorly explored.*

Intracranial recordings in neurosurgical subjects with a large number of electrodes spread over a relatively large surface of the cortical surface, a method known as electrocorticography (ECoG) (18) offers a new opportunity for acquiring fast temporal information from precisely localizable sources of signal. This method offers millisecond temporal resolution and millimeter anatomical precision - in the subject’s own native brain space. Unlike the uniform spatial coverage of imaging methods, intracranial EEG relies unfortunately on sampling from a limited number of implanted areas and leaves behind regions outside the coverage zones. While this introduces the problem of limited anatomical sampling, it may provide sufficient coverage for recording simultaneously from many sites within each individual brain in order to explore the spatiotemporal dynamics of activity across different cortical sites within each individual brain. In addition, the intracranial approach also allows delivering electrical pulses to discrete neuronal populations while causal changes in the subjective experience of the participant can be probed.

Using simultaneous recording across a relatively large area of the human brain one could test the hypothesis that *face information is first processed within the most face-selective sites that are anatomically discrete and localizable within individual brains and anatomically consistent across subjects, and that the information is then distributed to less selective sites*.

While recent intracranial recordings and stimulation studies (19-24), including our own (25-28), have addressed the neurophysiological underpinnings of face processing in the human brain, to our knowledge, these studies have yet to address the notion of anatomical selectivity and temporal distribution of face information using a multiprong approach (i.e., univariate and multivariate methods of recording and causal probing with electrical stimulation). The current study was designed to combine these methods to test our proposed alternative hypothesis.

We like to emphasize here that the aim of the study was not to decipher the complex computational code of face perception in the human brain. Studies in primate brains are perhaps much better suited for that purpose. The main purpose of the study was simply twofold: 1) to explore if regions of the temporal cortex that are selectively responsive to face stimuli are engaged earlier in time than those that are not face-selective, and 2) whether the stimulation of selective or non-selective ones cause the same effect in conscious viewing of faces. As one can imagine, these two questions could not be addressed with imaging methods because of their limited temporal resolution, or with single cell recordings because doing singe cell recordings from tens of cortical sites is not ethically or clinically possible in the human brain and or feasible even in the primate brain. Moreover, question 2 pertains changes in the conscious and subjective processing of faces and as such we could not do the study in animals since they cannot report such changes.

## RESULTS

We recruited 8 neurosurgical patients implanted with ECoG electrodes as part of their presurgical invasive evaluation for medication-resistant focal epilepsy. Subjects had unilateral electrode implantation in the right (5 subjects) or left (3 subjects) hemisphere (Table S1). Electrodes across all subjects (n=357) provided suitable coverage over the ventral and lateral TC.

We recorded from each of the 357 implanted electrodes with high temporal resolution (>1000 samples per second) while the subjects performed a visual task in which they viewed images of faces (human, mammal, bird, marine), and non-faces, including bodies without faces (same 4 categories), limbs (human), objects and places. They were instructed to press a button when a red hashtag sign appeared (Figure 1).

**Figure 1:**
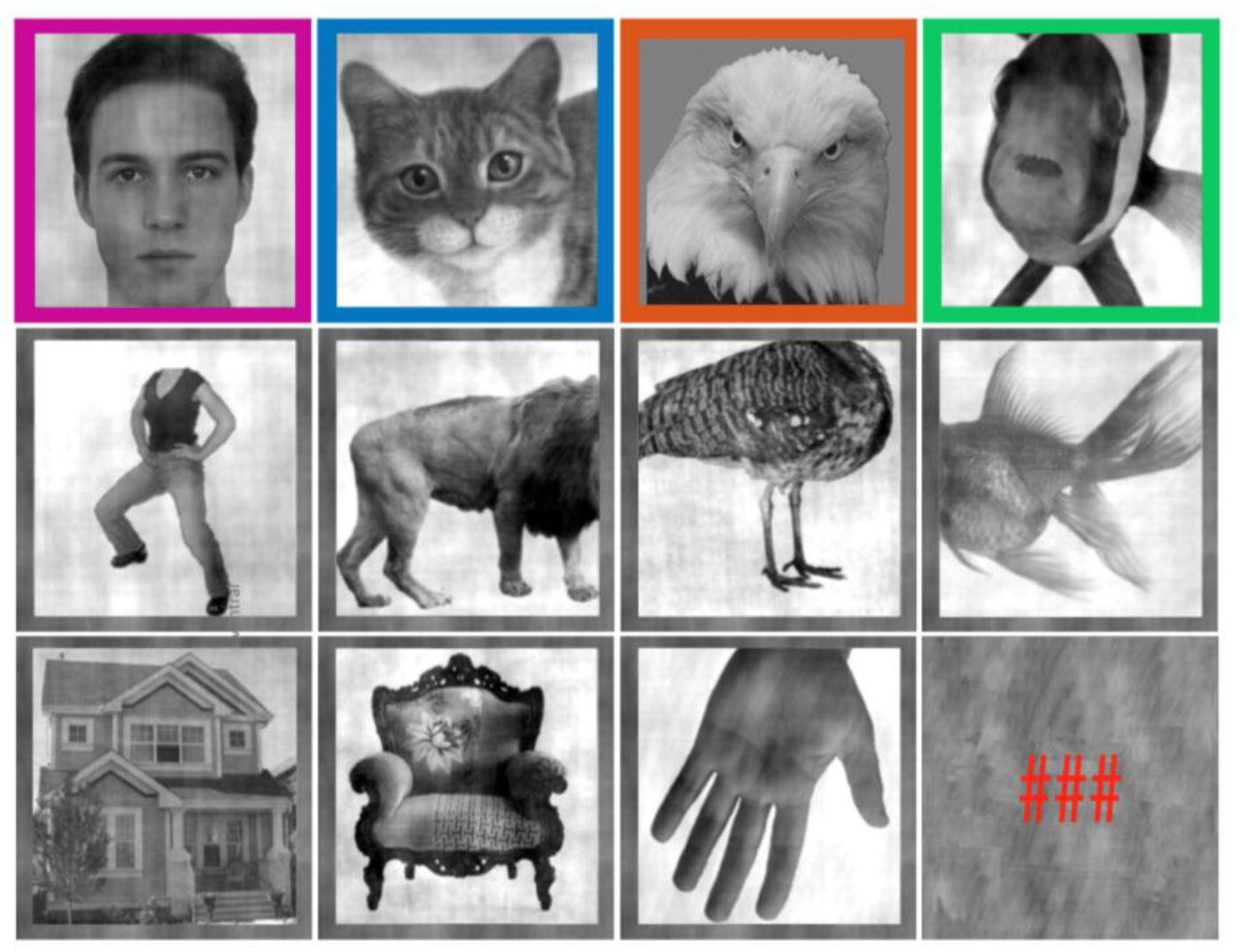
Experimental stimuli. Experimental task conditions with one exemplar of each image category displayed. The categories include human face, body and limb, mammal face and body, bird face and body, marine face and body, place and object. The subject was instructed to press a key when the target stimulus was presented (i.e. red hashtag sign). Low-level features were not different across categories. Four categories of face stimuli are shown here in different colors, and all non-face stimuli in gray frames only to correspond to the same colors used to denote these categories in the next figures.

We quantified the face information (i.e. the ability to discriminate faces from non-faces stimuli) in narrow bands of frequencies (θ: 4-7Hz, α: 8-12Hz, β_1_: 13-29Hz, β_2_: 30-39Hz and γ: 40-69Hz) as well as High-Frequency Broadband (HFB, 70-177 Hz) signal using univariate tests and a Multiple Kernel Learning (MKL)(29) method configured for intracranial EEG signals (30). Compared to the power of any other frequency band, the signal in the HFB showed the strongest univariate and multivariate changes in response to faces relative to non-faces (Figure 1a and Table S2). Given this finding, we focused further analyses on the HFB power and its profile of response across anatomical sites and task conditions.

We are mindful that the richness of the intracranial electrophysiological signal could have been explored by analyzing the power or phase of slower frequencies, or their coupling with higher frequencies(31). However, the HFB signal is well suited for the purpose of testing our predictions not only because of our MKL findings (Figure 1b and Table S2), but also because of the large body of evidence from other human(32-37) and non-human(38-43) studies (as summarized in (18)) that have confirmed HFB power as a reliable correlate of hemodynamic signal and averaged single and multi-unit activity of a population of neurons in a given cortical site. More importantly, HFB has a more precise anatomical source (i.e., micrometers around the recording electrode) compared to lower frequencies (several millimeters)(18). While the HFB signal (similar to BOLD signal) provides a suitable marker for the engagement of a given cortical site in a given function and as such, the time of onset and the power of HFB provide valuable information about the time and level of engagement of a *population* of neurons within tens or hundreds of micrometers of the recording electrode (41, 42).

We remind the reader that the purpose of the study was not to decipher the computations that are involved in each region of the brain during face perception. We simply wanted to identify the timing and level of cortical engagement and compare it across tens of different recording sites. As mentioned above, it should be noted that, compared to slower oscillations, the signal in the HFB showed the strongest univariate and multivariate changes in response to faces relative to non-faces.

Using HFB activity, we found a heterogeneous profile of responses across recordings sites (Table S3). Of the recorded sites (n=357), 53.22% (n=190) had significant responses to at least one category of stimuli relative to baseline (i.e., *“active”* sites). 13.45% recording sites (n=48) showed selective activations to human faces (i.e., *“human face selective”* sites) compared to any other stimuli (Figure 2a). Only 10.64% of the recording sites (n=38) showed face selective responses (comparing all sub-categories of faces to all non-faces; i.e., *“face selective” sites).* The (human) face selective sites were clustered in the fusiform gyrus or lateral occipital gyrus. Interestingly, While there was clear overlap between the “face selective” and “human face selective” sites, a few sites (17 out of 357 sites) showed selective responses to human faces while lacking significant responses to other face stimuli (Figs S1 and S2). In further analyses, we refer to *“task active*” sites as the sites that were assessed as “active” and that are not face or human face selective.

**Figure 2:**
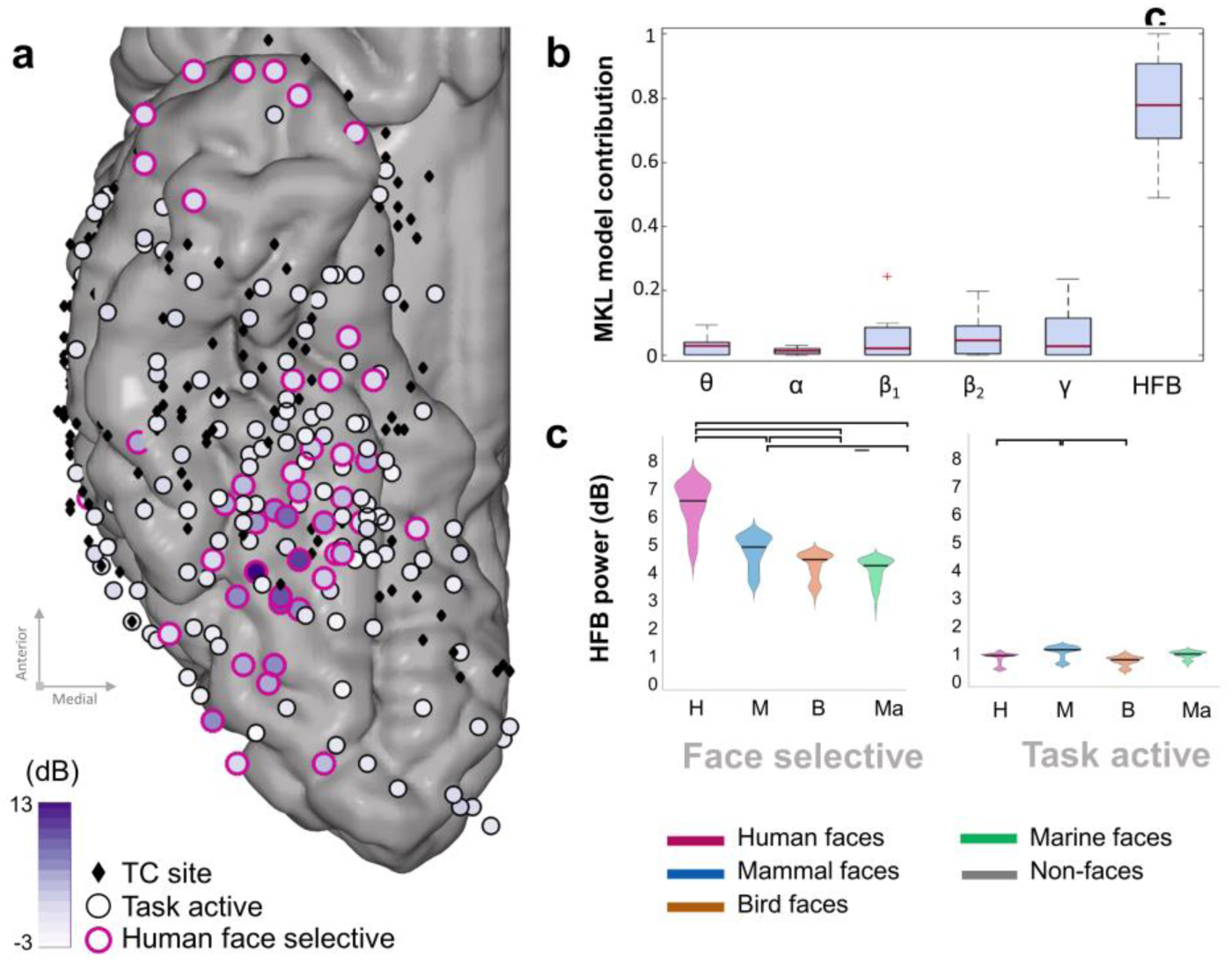
Effect of species on face coding. **(a)** Task active and human face selective sites across subjects in HFB. Among the 357 included TC sites (represented by black diamonds), 190 were task active (represented by circles) as defined by permutation tests (i.e. presenting a significant response to at least one category). The difference between their response to human faces and non faces stimuli are displayed as a color coded fill. 48 task active sites were identified as selective for human faces (represented with a pink contour) **(b)** HFB activity has the highest contribution in the discrimination between human faces and non-faces, within and across subjects. The results of the MKL model are plotted as box plots of the frequency band contributions to the model across the eight subjects, with the median represented by a red line. **(c)** HFB amplitude averaged within the [150 500]ms time window after onset for each of the 4 subcategories, at face selective (left) and task active sites (right). Coloring and initials represent the face subcategories: Human (H/pink), Mammal (M/blue), Bird (B/orange) and Marine (Ma/green). Given the size of the recording electrodes, and that the HFB represents the averaged neuronal population responses, it is likely that the electrodes labeled to be face selective sites recorded the activity of not only the face selective population of neurons but also their adjacent non-selective populations. For face selective sites, significant differences can be found (paired permutation tests, displayed by a black bracket) between the HFB responses to human faces and the mammal, bird and marine faces, as well as between the mammal and the bird and marine faces. There is no significant difference between the responses to bird and marine faces. For task active sites (i.e. active but not face selective), significant differences can be found between the bird faces and the human and mammal faces.

To explore whether biological similarity of the face stimuli (humans and mammals versus birds or marine) influences the neural responses in the face selective sites, we compared responses to faces of different categories. This analysis showed clearly that human faces induced the strongest activations on face selective sites (median=5.21dB, n=1579) followed by mammal faces (median=4.06dB, n=1609), bird faces (median=3.48dB, n=1666) and marine faces (median=3.48dB, n=1706, Figure 2c left). All pairs of face subcategories showed a significant difference in HFB power (permutation test, p<0.05 after Bonferroni correction), except for the comparison of bird and marine faces (human-mammal: p<0.0001, human-bird: p<0.0001, human-marine: p<0.0001, mammal-bird: p=0.0012, mammal-marine: p=0<0.0001, bird-marine: p=0.9896). In comparison, the amplitude of responses to subcategories of faces in the task active sites showed lower HFB power for the face categories (human faces=0.73dB, n=5761, mammal faces=0.91dB, n=5497, bird faces=0.67dB, n=5837, marine faces=0.79dB, n=5808, Figure 2c right). Only mammal faces elicited significantly higher responses than did human and bird faces (mammal-human: p<0.001, mammal-bird: p<0.0001). Other pairwise comparisons did not show differences in terms of HFB amplitude across the four subcategories of faces (p>0.05, after FDR correction). Please note that these univariate results were not driven by physical differences in the stimuli (Supplementary S4).

Given that the HFB responses to human faces had the highest signal to noise ratio, further analyses focused on the processing of human face stimuli.

To address the earlier imaging reports of distributed face processing (17), we investigated decoding of face information across human face and task active sites. To this end, we referred to a framework (45) which allows inferring causal relationships between the stimuli and the observed EEG activity in encoding (e.g. univariate analyses) and decoding (e.g. machine learning based modeling) settings. This is of interest as decoding performance or machine learning model contribution cannot be directly *causally* related to the source of the signal. Instead of referring to decoding performance, this framework assesses the ‘relevance’ of both face and task active sites in discriminating between human face and non-face epochs. A feature is assessed as ‘relevant’ in decoding settings if, when removed, the performance of the model is significantly affected. This is similar to the procedure used in a recent publication(46), in which the face information shared across regions of interest was taken into account. In encoding settings, features are relevant if they display significant responses/contrasts in a univariate test. Features assessed as (respectively not) *relevant* in both encoding and decoding settings are (respectively not directly) causally related to the stimuli. In our work, we ran three machine learning schemes discriminating between human face and non-face stimuli, using signals from different sets of sites. The results from all models are displayed in Table S5 and Figure 3a. For each subject, Model I incorporated data from all TC sites (results displayed by dark grey triangles on Figure 3a). For all subjects, Model I was able to significantly discriminate between human faces and non-faces (permutation test p<0.05, FDR corrected for number of subjects). Model II used the same classification but excluding the face selective sites (red triangles in Figure 3a). Significant discrimination between faces and non-faces was only observed in 2 subjects out of 8 (permutation test, p<0.05 FDR corrected for number of subjects). In these 2 subjects, the significant decoding accuracy suggests that at least one site contains information about the discrimination at hand. This could however be related to selective responses to non-face stimuli, as we had not excluded sites selective to one or more subcategories of non-face stimuli. Comparing Models I and II revealed that there is a significant decrease in accuracy when excluding face selective sites from the classification across subjects (Wilcoxon signed rank test, p=0.0078, n=8). This result suggests that face sites are *relevant* in decoding settings for face processing (45).To test whether random subsets of task active sites are relevant in decoding, we built 499 models that included the same number of sites as Model II by randomly selecting task active and face selective sites, but including at least one face selective site. This means that across the 499 models (referred to as ‘random set’ models), the proportion of face selective sites varied, but was non-null. The performance of these models is represented by colored dots on Figure 3a, their color displaying the proportion of face sites included in the model (dark blue is 1 face site, light green is all face sites). Across subjects, removing random sets of task active and face selective sites did not affect the model performance significantly (Wilcoxon signed-rank, p=0.5781), nor did it within subjects for 7 out of 8 (Table S5). For Subject 2, a multimodal distribution is observed reflecting whether or not a specific face selective site is included in the model (leading to accuracies higher than 79%) or not (accuracy lower than 70%, 83 models out of 499). There is a significant difference in model performance when this site is included or excluded from the model (p<0.0001). These results suggest that task active sites were *not relevant* for the decoding model. It is further supported by the fact that non-significant classification accuracy is observed in Model II for 6 out of 8 subjects. It is also interesting to note that there is a significant relationship between the proportion of face sites included in the analysis (i.e. from one face site to all sites, randomly selected in the 499 models) and the model performance for 6 out of 7 subjects (Table S5, n= 499). This result could arise from two scenarios (or a combination of the two): either face information on face sites is not redundant, i.e. each face site brings unique face information and/or including more sites leads to better signal-to-noise ratio of the face pattern. In both cases, this result suggests that the pattern can be more easily identified when more human face sites are included.

**Figure 3:**
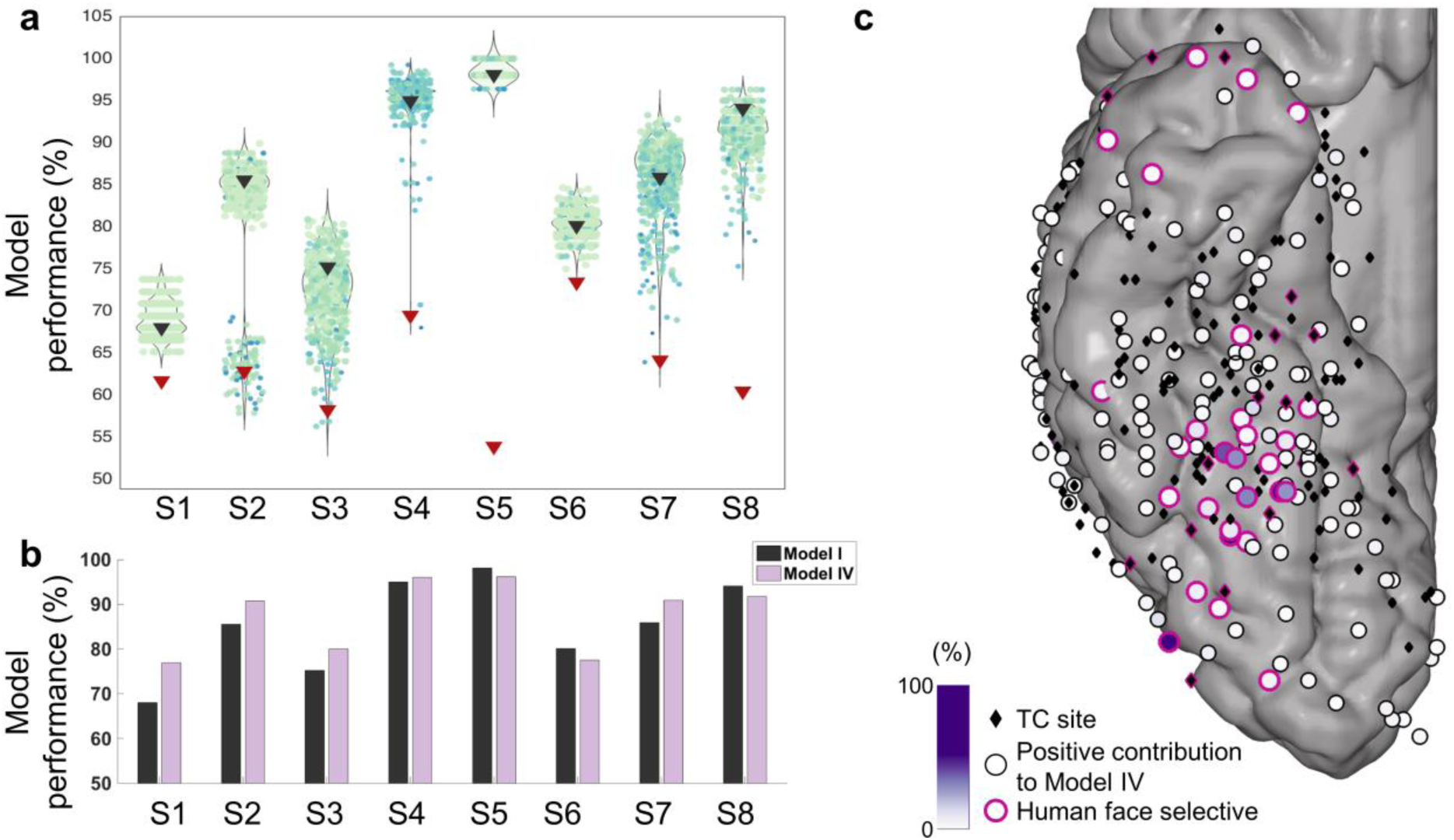
Relevance of TC sites in decoding settings. (**a**) Dark grey triangles represent the performance of Model I, i.e. including all TC sites. Performance of Model II is represented by red triangles. Each colored dot represent one of the ‘random set’ models (499 models per subject), their color representing the proportion of face sites were included in the model (dark blue is 1 face site, light green is all face sites). Violin plots represent the distribution of the ‘random set’ model performances compared to Model I and Model II. (**b**) Sparse models perform as well as or better than distributed models. Bar plot representing the balanced accuracy for Model I compared to the model accuracy of sparse Model IV, for each subject. (**c**) Site contributions to the sparse model, plotted across subjects. The contribution of the site (in %) is represented by a color-coded fill. Black diamonds have a perfectly null contribution to the model while circles had a positive contribution to the model. Sites assessed as human face selective by the univariate analysis are highlighted by a pink rim.

The degree of inter-subject variability (both for Model II and the ‘random set’ model) could not be explained by the number of trials (ρ(Model II - #trials) = 0.5073, p=0.1994, n=8; ρ(median random sets - #trials) = 0.0068, p=0.9872, n=8) or of sites included in the analysis (ρ(Model II - #sites) = −0.1221, p=0.7734, n=8; ρ(median random sets - #sites) = 0.00156, p=0.9708, n=8). The inter-subject variability, however, could be explained by various factors such as signal-to-noise ratio, amount of correlated noise or placement of the electrodes, which are complex to quantify. See Materials S6 for a discussion on circular analysis and its effect on the presented results.

To explore the results further, we performed an additional machine learning model to assess the anatomical distribution of face processing in the human TC. In this analysis, the classification is the same as for Model I, except that the considered algorithm enforces sparsity at the site level, i.e. it automatically selects a subset of sites to perform the classification. Comparing sparse and non-sparse modeling techniques is a common machine learning strategy to investigate data properties: SVM (i.e. Model I) assumes that the information is fully distributed across features, by construction. If this assumption is correct, it should perform better than a sparse model. By contrast, if the sparse model performs better than Model I, it suggests that the information contained in the selected subset of sites is sufficient to perform the classification and that the non-selected features might increase the noise (i.e. not bring relevant information). As in previous publications(30, 47), we used the simpleMKL algorithm (29) to perform the sparse modeling. Please note that the basis algorithm for the simple MKL is an SVM, hence the effect of implementation on the results is limited (29). The results are displayed in Figure 3b and Table S7. In all subjects, the sparse model performs either significantly better than, or equally well as, the distributed SVM model (Wilcoxon signed-rank test, p=0.0312, n=5). In Subjects 5, 6 and 8, the difference in model performance between Model I (SVM) and Model IV (sparse MKL) is not significant (Wilcoxon signed-rank test, p=0.1250, n=3). Of note, in two subjects the accuracy of the model was over 90%, which left little room for improvement.

In addition, the sparse model weight maps significantly overlapped with the univariate maps of human face selectivity across subjects (ρ = 0.4708, p<0.0001, Figure 3c) and within subjects for 7 out of 8 subjects (except Subject 8, Table S7 and Figure S3). This result shows that MKL relies heavily on face selective sites for the classification. For Subject 8, the classifier seems to rely on non-face information, as the site with highest contribution contains signals larger for all non-face categories than for human faces (pooled non-face>face: p<0.05).

Our univariate and multivariate results display that face sites are ‘relevant’ for human face processing. However, we also showed that including more face sites to the model increases model performance (random set models). We then explored the relationship between selectivity, anatomy and timing of the HFB responses on face and task active sites during human face processing.

While the face sites displayed significant responses to human face stimuli, the selectivity of their responses decreased from posterior to anterior sites as demonstrated by correlating the MNI y-coordinate with selectivity (ρ= −0.6191, p= 3.51e-06, n= 47). For task active (i.e. not face selective) sites, selectivity to human faces increased from posterior to anterior sites (ρ= 0.2744, p= 0.0011, n= 139).

Our fast event-related paradigm combined with simultaneous recordings across selective and non-selective sites with high sampling rate allowed us to compare the latency of neuronal population responses to faces within the first 500ms of stimulus presentation. We measured the Response Onset Latency (ROL) to human face stimuli across human face selective sites and task active sites based on unsmoothed, normalized signals (Figure 4). We found a clear posterior-to-anterior lag in time of onset of HFB responses to both human face selective and task active sites. More posterior human face selective neuronal populations responded significantly earlier than did more anterior ones (i.e. ROL values across human face selective sites were significantly correlated with the y-coordinate of the corresponding site, ρ= 0.4692, p=0.0034, n= 37). Interestingly, the posterior-to-anterior lag was also significant for task active sites (ρ= 0.6567, p=7.1995e-07, n= 46). It is noteworthy that the ROL technique identifies task active sites that respond non-selectively to human faces (see methods) to assess a latency value. More importantly, the group level findings presented here were also present at the individual subject level for human face selective sites, when calculating the ROL for each site relative to the most human face selective site within subject (Figure 4b,c): ρ(ROL-y) = 0.7593, p=1.7911e-06, n=29 and ρ(selectivity-y) = −0.5680, p=0.0013, n=29. In other words, the posterior to anterior ROL gradient was not driven by one or a few subjects who happened to have coverage over a specific region of the TC. For task active sites (Figure S4a-b), the correlation between ROL and y-coordinate was also significant: ρ = 0.6756, p=2.5957e-7, n=46. On the other hand, there was no significant correlation between anatomical position and selectivity when compared to the most face selective site: ρ = −0.2221, p=0.1380, n=46. There is no significant difference in ROL values between human face selective and task active sites (p=0.9452, Wilcoxon rank sum test).

**Figure 4:**
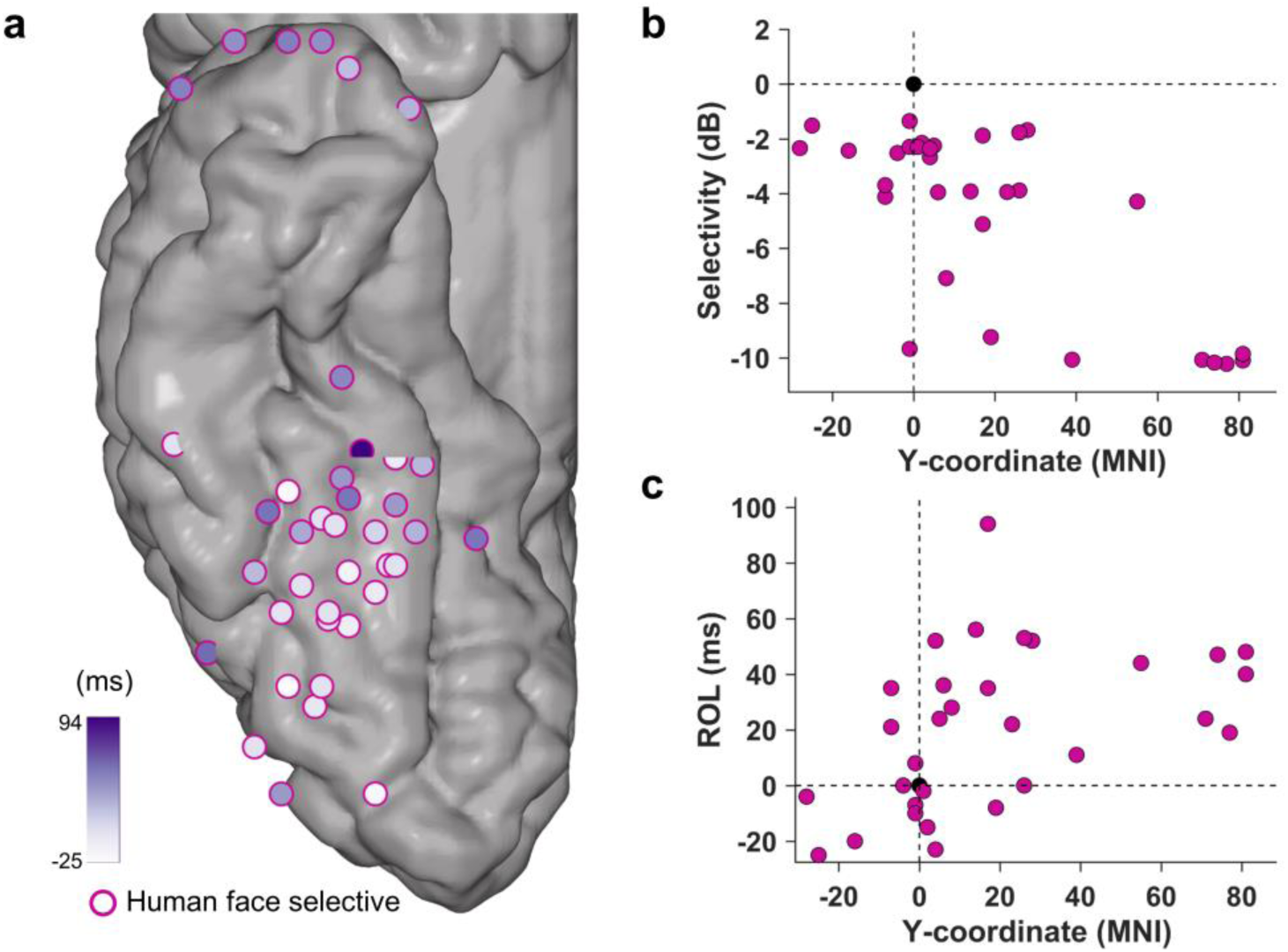
Temporal distribution of human face information. **(a)** Response Onset Latency (ROL) over human face selective sites for the human face category, as represented by a purple color scale fill on the MNI cortex with the best (i.e. most human face selective) site in each individual chosen as the point of reference. **(b)** Selectivity in each subject, when compared to MNI y-coordinate with the best site (i.e. most human face selective) in each individual chosen as the point of reference. The best site is represented as a black circle, at the crossing of the 2 axes. Selectivity is related to anatomical position. **(c)** ROL in each subject, when compared to MNI y-coordinate with the best site in each individual chosen as the point of reference. Latency is related to anatomical location of the electrodes.

Importantly, the effect presented here does not result from biases to signal amplitude or slope in our ROL method (supplementary information S8 and Figure S5).

Lastly, we explored the effect of electrical perturbation of face selective sites on subjective processing of faces. For this, we hypothesized that the stimulation of more posterior sites (with most selective and earliest responses) would cause more salient effects than the anterior sites. This hypothesis has not been addressed in prior electrical stimulation studies showing distortion in conscious viewing of faces (26-28, 48, 49) or naming of famous faces (50, 51). Subjects were instructed to view a human face at the bedside while we performed active or sham (zero current) stimulations of selective, task active or non-responsive sites. Subjects then reported whether the face remained the same or was distorted. We emphasize that this anecdotal report departs from the well-controlled experimental procedures that could be performed in non-human primates (14, 52). However, the limitation of performing the procedure in a clinical setting and in patient populations precluded such experimentally rigorous studies in our subjects.

The complete report of the face-related stimulations, verbal prompts from the doctor, and the patient’s verbal and non-verbal responses can be found in Supplementary S9. Distortion of human face perception was reported only with real (and not sham) stimulation of some, but not all, face selective sites, with stimulation of more posterior sites causing the perceptual perturbations (Supplementary S9). Stimulation of only a few face selective sites in Subjects 4, 5, 6, 7, and 8 resulted in the distortion of human face perception, while stimulation of other selective or task active sites in the same individuals caused no change (Figure 5). Given the anatomical locations of these sites, the general trend indicated that stimulation of more posterior fusiform sites causes distortion of face perception while the stimulation of face selective electrodes located in relatively more anterior areas tends not to yield the same change in face perception.

**Figure 5:**
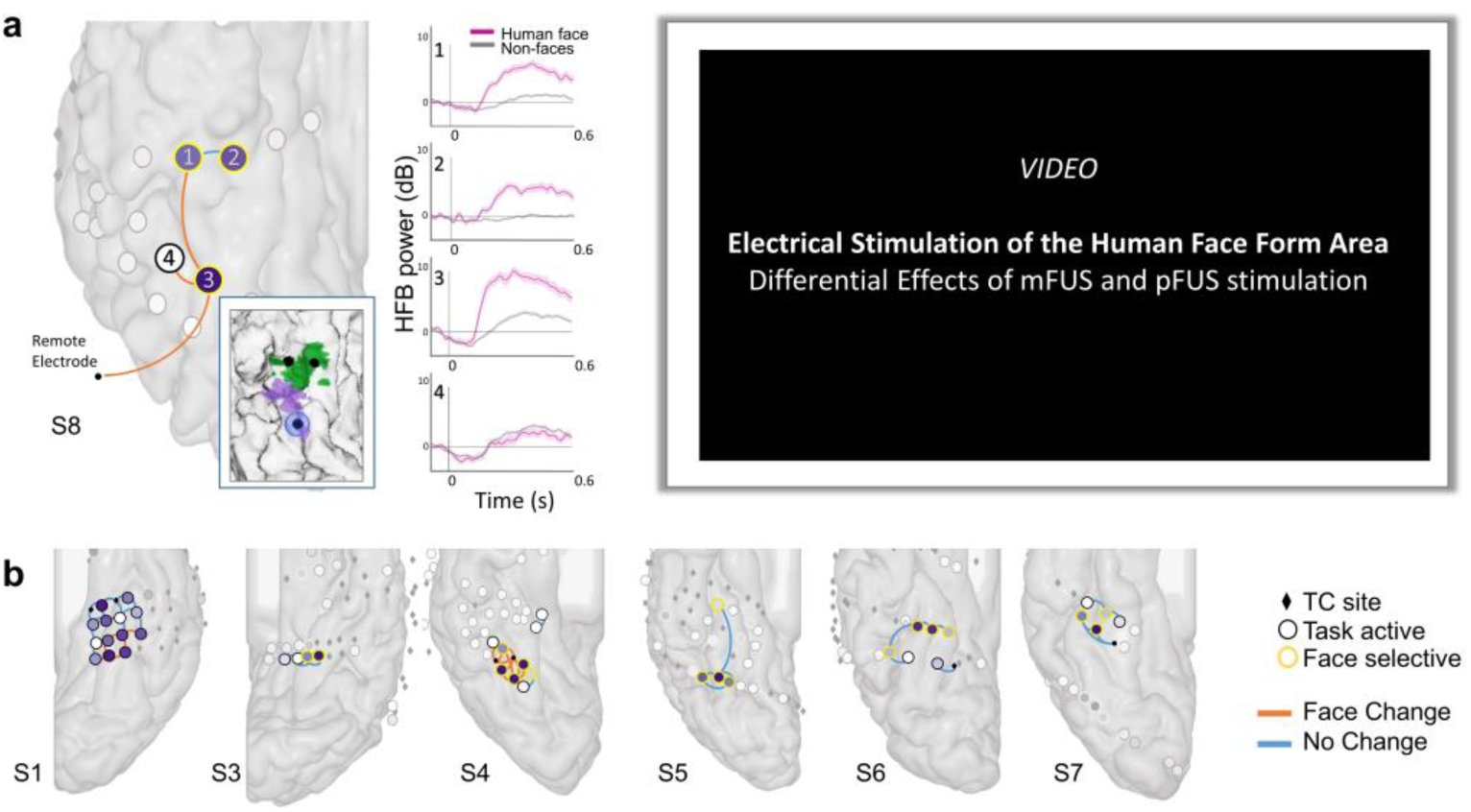
Electrical Brain Stimulation. The effect of bipolar stimulation of two electrodes is shown in orange lines (face distortion) or blue (no face distortion). (**a)** Sites 1, 2, and 3 are located in the fusiform gyrus and given their anatomical coordinates represent the fusiform face areas (FFA). In the box, the location of electrode 1, 2, and 3 are shown on top of a functional mapping of mFUS (green) and pFUS (purple). The estimated cortical area affected by the stimulation on electrode 3 was calculated using the relationship between the estimated charge per trial and the cortical area affected (Winawer & Parvizi 2016) and is shown with a blue circle around it. Site #3 is the site whose stimulation caused distortion of faces when it was stimulated in pairs with sites #1 (another face elective site), nearby site #4 (task active site) or a remote reference electrode (in the motor cortex). Middle panel shows the HFB responses to human faces (pink) and non-faces (grey) of the sites that were stimulated. The right panel has an imbedded video of subject S8’s verbal responses after being stimulated. The patient has consented to the use of his video in this publication. (Dear Reviewer, due to the large size of the video, we were unable to submit the video along with this manuscript. Please download it from https://www.dropbox.com/s/kdgmex01c4sn3g2/Figure5Video.mp4?dl=0. (**b)**. Localization of stimulated electrodes on Subjects 1, 3, 4, 5, 6, and 7 are depicted using the same convention as in Figure S2.

In one subject (Subject 8), we probed the effect of stimulations across two patches of the fusiform face area (FFA, numbered as 1, 2 and 3 in Figure 5a). Using bipolar (adjacent pair of sites stimulated together) or unipolar (site was paired with a distant reference) stimulations, we elicited distortions in seeing faces when the subject looked at his own face in the mirror, looked at a cartoon face drawn on a piece of paper, and focused on the eyes or the lips of the face. He also reported induced perceptions of a face during electrical stimulation while his eyes were closed. The subject’s verbal reports after stimulation include “one side of the face changed”; “facial features [turned] into a cartoon”; one eye “became someone else’s”; “face wiggled a little bit”; and “face looked familiar” (Video in Figure 5). More importantly, the effects were observed only when site #3 was stimulated. To explore the anatomical location of site #3, we localized the posterior fusiform face area (pFUS) and the medial fusiform face area (mFUS) onto Subject 8’s native neuroanatomical space. Using methods described in (53) the calculated field of electrical stimulation of site #3 is precisely localized in the pFUS (Figure 5a). The other face selective sites (#1 and #2), located in medial fusiform area (mFUS) failed to cause any distortions even though they were only 1cm away from site #3. In our previous stimulation report (28), we stimulated both mFUS and pFUS in a bipolar manner. This study is, to our knowledge, the first to differentiate the effect of pFUS versus mFUS stimulation using unipolar stimulation.

## DISCUSSION

Our study addresses the spatiotemporal distribution of face information based on univariate measures, timing analysis, machine learning based modeling, and direct cortical stimulation. Using this multi-pronged approach and by leveraging the temporal resolution of the ECoG method, we confirmed that the majority of recording sites in the human TC do not show any significant change of activity in response to visual presentation of faces while a minority of sites respond non-selectively to all categories of visual stimuli and only fewer sites, which are anatomically consistent across individual, respond selectively to face stimuli.

Our results suggest that removing face selective sites from machine learning based classification significantly drops the decoding accuracy of the model. In addition, task active sites were not assessed as *relevant* when discriminating human face from non-face stimuli. This was further supported by an improvement in model performance when considering a sparse approach. This is in disagreement with the previous imaging findings suggesting that face information can be decoded from non-selective sites. This discrepancy may be due to the differences in the number of recording sites, the time scales considered, and the methodology used. For instance, the number of anatomical samples in each subject is limited with the ECoG method, which can reduce the power of the algorithm in detecting weak, distributed patterns. To maximize the detection of small effects, we used Support Vector Machine classifiers, which are known for their ability to detect subtle, distributed patterns and we focused our analysis on the high signal-to-noise ratio of HFB responses induced by human face versus non-face stimuli. While we could not relate the number of sites to model performance (see Results), it is possible that the signal in task active sites is too weak to be detected across a few tens of anatomical samples in each subject. On the other hand, the novelty of our data is in part due to the high temporal resolution of ECoG that enabled us to measure the fast temporal dynamics of face processing in the human brain. For instance, our electrophysiological analysis relied on the responses elicited within 500ms of stimuli presented in an event related paradigm for 300ms and with an inter-stimulus interval of 400ms. This is in stark contrast to some of the classic neuroimaging studies whose temporal window included >10 seconds of signal processing (e.g., 24 or 16s long blocks of visual stimuli for each category (17)).

In a recent study(46), a multivariate analysis was performed on fMRI data to discriminate between different visual categories (faces, fruits/vegetables, letters and vehicles) in seven pre-defined regions of interest (ROI). The analyses were performed on each ROI separately, then on all regions together except one. The latter aims to display the unique contribution of each ROI to the discrimination, i.e. any shared information across ROIs will be taken into account in this scheme. The authors concluded that face information is *both* localized and distributed, but due to limited temporal resolution of the imaging method, authors could not confirm the hypothesis of *distribution in time*, as we have done here.

Another methodological caveat that needs to be considered is that the task active areas in humans may be variable in their relative size and primary cytoarchitectonic composition (54). Therefore, hubs of activity in posterior to anterior TC may have different sizes or shapes of physiological responses or locations on the surface of gyri vs. depth of sulci. As such, the results we report here could represent an idiosyncrasy of our intracranial EEG method. However, it is still noteworthy that responses to faces were significantly faster in the posterior sites than the anterior face selective sites. By using ECoG recordings, our results clearly confirm our overarching hypothesis that face information is anatomically localized but temporally distributed. However, we note that our study was not designed to determine the path of information flow, namely whether the face information in the non-selective sites comes directly from posterior face selective sites.

Additionally, our findings support the causal link between some, but not all, face selective sites of the fusiform gyrus and conscious processing of faces as suggested previously by us (26-28) and others (49, 50), but clearly demonstrate that stimulation of different face selective sites leads to different effects on conscious processing of faces. Moreover, our work suggests that the perturbation of the pFUS is more important for conscious face processing than the stimulation of mFUS. This is an intriguing finding that needs to be verified in a larger sample of subjects. As demonstrated in non-human primates (14, 52), and also in the human lateral occipital cortex (19, 22), we acknowledge that the face selective sites outside the fusiform gyrus might also play important and causal roles in face processing. A recent study in non-human primates clearly showed that the micro-stimulation of the very anterior face selective patch (area AM) severely distorted the monkey’s percept of facial identity, such that faces depicting the same identity appeared to depict different identities (52).

In closing, our data suggest that a few anatomically consistent sites play a crucial role in processing face information and that there is a time delay in their processing of the same information. However, our study does not suggest that the conscious perception of faces solely depends on the operation of these isolated patches of face selective sites. We acknowledge that different facets of face processing may occur in different patches of face responsive sites and that these neuronal patches are embedded in a larger network of visual and other association areas of the brain, and their function should not be seen in isolation from this wider integrated brain network (16, 55). Future studies with simultaneous recording across face selective and other association areas are needed to determine how different facets of face information is decoded in each of the face patches and how their function is embedded and relayed to the rest of the brain for serving human cognition and behavior.

## ACKNOWLEDGEMENTS

This work was supported by funding from the European Union’s Horizon 2020 research and innovation program under the Marie Sklodowska-Curie grant agreement No 654038 (DecoMP-ECoG) to JS; National Science Foundation Graduate Research Fellowship (DGE 1106400) to VR; National Institutes of Child Health and Human Development NRSA postdoctoral fellowship (1F32HD087028-01 from) to ALD; research grants from the US National Institute of Mental Health (1R01MH109954-01) to J.P. We are thankful to Dr. Kevin Weiner for generous help in creating Figure 5a; Dr. Larry Shuer and Dr. Hong Yu for performing the implantation procedures for clinical reasons; and to Harinder Kaur, Thi Pham, and Luda Schumacher who were the clinical EEG technologists providing support during the research recordings.

## Code availability statement

All analyses were performed in Matlab (www.mathworks.com). Pre-processing and univariate analyses were performed based on SPM (http://www.fil.ion.ucl.ac.uk/spm/) and in-house routines available at https://github.com/LBCN-Stanford/Preprocessing_pipeline. ROL in-house codes are available on Github at https://github.com/LBCN-Stanford/. Multivariate analyses were performed using a development version of PRoNTo (30, 56). This code will be released as PRoNTo v3 and be available at https://github.com/JessicaSchrouff. The code to build semi-simulated data is available at https://github.com/JessicaSchrouff/Simulated_ECoG, along with the rest data from subject S1 used to generate the noise structure.

## METHODS

### Demographics and recordings

Eight subjects (six males, two females, aged between 23 and 68 years) were implanted with intracranial electrodes to localize the source of drug-resistant seizures. The procedure was approved by the Stanford Institutional Review Board (IRB) and the subjects provided written informed consent to participate in the study. The location of the grids was determined by clinical needs (Figure S1, three left hemisphere implantations, five right). Data were obtained at 1525.88 Hz through a 128-channel recording system (Tucker Davis Technologies, http://www.tdt.com) for the first seven subjects while a Nihon Kohden Technology system with simultaneous video monitoring was used to perform 1 kHz recordings in subject S8. Each electrode was a platinum plate, either 2.3mm or 1.15mm in diameter (exposed recording area) with center-to-center spacing of 4–10 mm between adjacent electrodes on the grid or strip. Electrodes containing artifacts or pathological activity were discarded from further analyses.

### Anatomical localization of electrodes

Structural MRIs were acquired with a GE 3-Tesla Sigma scanner at Stanford University equipped with a head coil of a T1-weighted SPGR pulse sequence was AC-PC aligned and was resampled to 1mm isotopic voxels, then segmented to separate gray and white matter. Post-implantation CT images were aligned to the pre-op MRI anatomical brain volume (57). Electrodes were visualized on the subject’s own brain volume and reconstructed onto a 3D cortical surface allowing for accurate anatomical localization of electrodes. The electrode positions were also transposed into the MNI space and displayed on a MNI cortex file for visualization of results across subjects.

### Experimental paradigm

The experiment was administered using psychtoolbox (http://psychtoolbox.org/) running on Mac OSX. The laptop was placed ∼70 cm from the subject’s eyes at chest level. Screen resolution was 1280×800. Each image was subtended 5 visual degrees at its longest dimension. Each subject underwent a visual task during which images of different categories were presented at the center of the screen for 300ms, with an ISI of 400ms (see Figure 1 for representative examples). The categories included human face, human body, mammal face, mammal body, bird face, bird body, marine face, marine body, human limbs, object and place. The image backgrounds were phase-scrambled at 3% in order to reduce visual artifact. The visual dimensions of the image plus its scrambled background were 11.10cm x 11.10cm, and the visual angle was 9 degrees. Each category comprised 25 images, presented twice. This hence leads to 50 stimuli per category and 550 visual stimuli in total. During image presentation, the subject was asked to press a key (‘press 1’) when the pattern ‘###’ appeared in red at the center of the screen (further referred to as a ‘Response Block’). The onset of each stimulus was recorded by a photodiode signal generated by a luminance change in the display at image onset.

### Signal Preprocessing

All preprocessing steps were performed using Matlab (<u>The</u> MathWorks, Inc., Natick, <u>Massachusetts</u>) and the SPM (www.fil.ion.ucl.ac.uk/spm) toolbox in custom routines (https://github.com/LBCN-Stanford/Preprocessing_pipeline). The data was first down-sampled to 1000 Hz and filtered for power-line noise (band-stop between 57 and 63Hz) and harmonics (117 to 123 Hz, 177 to 183 Hz). Sites underwent an automatic quality assessment: sites with variances 5 times larger or smaller than the average variance across all sites were labeled as pathological and excluded. Sites with 3 times more ‘jumps’ (defined as changes in the signal derivative larger than 100 μV) than the average across sites were considered as spiky and excluded. The signal was then re-referenced to the average of the signal over all selected sites. Each event was extracted (i.e., epoched) in the −200 to 700ms time window around its onset and baseline correction was performed (using the [-200 to 0]ms time window around onset as baseline). Events were marked as artifacts if they contained spikes of >100μV, and were discarded from further analyses. A time-frequency decomposition was then computed using a 7-cycle Morlet wavelet, with frequencies ranging from 70 to 177Hz (steps of 1Hz, avoiding discarded frequencies from Notch filtering). A similar 5-cycle Morlet wavelet time-frequency decomposition was performed for frequencies ranging from 1 to 69Hz. The power in each frequency and time bin was rescaled using the log-power of the [-100 0] ms window around stimulus onset. Six frequency bands were considered in this work: θ (4-7Hz), α (8-12Hz), β1 (13-29Hz), β2 (30-39Hz), γ (40-69Hz), and High Frequency Broadband (HFB, 70-177Hz). The signal was finally averaged across frequency bins within each band considered and smoothed with a 50ms width Gaussian window. The HFB power in the [-100 600] ms window around stimulus onset was considered for further analysis (see supplementary S2 for justification).

### Relevance of TC sites in encoding

For each subject, sites were defined as ‘active’ if they displayed a significant HFB response in the [150 500]ms after stimulus onset for at least one category, when compared to the event baseline ([-100 0]ms before onset). Paired non-parametric permutation tests (50,000 permutations) assessed the significance of the response (p<0.05, FDR corrected for the number of sites tested). Out of the active sites, ‘face selective’ sites were identified as sites where significantly higher responses were seen to the 4 face categories (human, mammal, bird, marine) pooled compared to all other stimuli pooled (non-parametric permutation tests, FDR corrected) in the time window [150 500]ms after onset. Finally, ‘human face selective’ sites displayed significantly higher responses to human faces compared to all non-faces stimuli pooled (non-parametric permutation tests, FDR corrected) in the [150 500]ms after onset window. Active sites that were not assessed as face selective or human face selective are further referred to as ‘task active’ sites. Sites assessed as (human) face selective were considered as *relevant* in encoding settings for face processing(45). These analyses were performed for the HFB signal but also for other frequency bands (see supplementary S2). In Supplementary S4, we investigate the influence of low-level image features on the univariate results. The four face subcategories were then compared on both the ‘face selective’ and ‘task active’ sites based on the amplitude of their HFB response averaged in the [150 500]ms time window after onset. Permutation tests assessed the significance of potential differences between subcategories (10,000 permutations, FDR corrected for the number of tests (n= 12, 6 binary comparisons for face sites and 6 for non-face sites)). Please note that there is no circularity in this analysis as the contrast to select sites (i.e. faces versus non-faces) is different from the effect investigated (i.e. human faces vs mammal faces vs bird faces vs marine faces).

### Frequency information for faces

For each frequency band, a univariate analysis assessed active, face selective and human face selective sites. In addition, a multiple kernel learning (MKL) (29) model assessed the contribution of each frequency band to the discrimination between human faces and non-faces (30). In this case, a linear kernel is built for each frequency band. Those kernels are then combined during the modeling step, based on a sparsity constraint. The model outputs a contribution for each kernel that can be interpreted as the weight of each frequency band in the classification. All modeling parameters were kept consistent with models I, II and IV (see ‘*Relevance of TC sites in decoding*’).

### Low-level image features

To ensure that low-level features in the stimulus images did not drive our results, we performed multiple control analyses.

### Stimulus features

First, spatial frequency power spectrum with rotational average was calculated for each stimulus and averaged across categories. Averagedspatial frequency power spectrums were compared using the one-way Anova and post-hoc analysis was performed using ‘Tukey-Kramer’ method. Power spectral analysis returned 153 values for each image, which were averaged within each category (n=153). We also computed the mean luminance across pixels in each category, pooling all non-faces together.

### Univariate analysis

we investigated the effect of mean luminance on the univariate results (i.e. on sites defined as ‘face’ selective). To this end, we plotted the histogram of mean luminance in the faces and in the non-faces category. We defined as low (resp. high) luminance faces, face stimuli with a mean luminance smaller (resp. larger) than 120 (threshold defined arbitrarily). The neural signals in each category was compared on all face sites (high luminance faces: n=45, human:15, mammal:12, bird:7, marine:11, low luminance faces: n= 55, human:10, mammal:13, bird:18, marine:14), using permutation tests.

### Relevance of TC sites in decoding

We then assessed the *relevance* of face and task active sets of sites in decoding settings. To this end, a machine learning model discriminating between human face epochs and non-face epochs was estimated based on three different site sets: (I) All TC sites included for analysis (referred to as the ‘TC’ model); (II) All TC sites, excluding the ones assessed as ‘human face selective’ by the univariate, permutation tests (referred to as the ‘TC-sign’ model); and (III) 499 random subsets of task active and face sites, including at least on face site. In scheme (III), further referred to as the ‘random sets’ model, the number of sites (i.e. features) included for modeling is identical as for Model (II). However, the proportion of face sites randomly varies, from 1 to all face sites. The different models aim at answering the following questions: (I) Is it possible to significantly discriminate human face from non-face trials in each subject? (II) Are face selective sites *relevant* for the discrimination between human faces and non-faces? I.e. do we observe a significant change in model performance when removing the set of face selective sites? Accessorily, is it still possible to significantly discriminate between human face and non-face trials? Or, on the contrary, is the information in those human face selective sites *necessary* for significant classification? (III) Are task active sites *relevant* for the discrimination between human face and non-face stimuli? I.e. is model performance significantly affected when removing random sets of task active sites? This analysis was conducted in PRoNTo version 3.0 (30, 56). The data considered focused on the [150 500]ms after stimulus onset. The ‘mammal body’, ‘bird body’, ‘marine body’, ‘human body’, ‘object’, ‘place’ and ‘limbs’ categories were pooled together to form the ‘non-face’ category. As this leads to imbalances in terms of the number of trials in each class (maximum 50 human faces compared to maximum 350 non-faces), epochs from the ‘non-face’ category were randomly subsampled to closely match the number of epochs in the ‘human face’ category. During this process, care was taken to include approximately the same number of epochs from each sub-category (e.g. 7 ‘mammal body’, 8 ‘bird body’, 7 ‘marine body’, 7 ‘human body’, 8 ‘object’, 7 ‘place’ and 7 ‘limbs’). A linear kernel matrix was built based on the data from all sites considered in the feature set (number of features: number of sites x 351 time points). This matrix corresponds to a similarity matrix between each pair of epochs (dot-product). The similarity matrix was then input into a Support Vector Machine classifier. It should be noted that SVM is an L2-norm regularized technique. This means that it does not assume or enforce a sparse distribution of the model weights. It should hence, in theory, be able to identify subtle, distributed patterns over the TC. Model performance was computed based on a 5-folds cross validation, i.e. 20% of the epochs were left out before training the model on the 80% remaining epochs (non-overlapping). The model was then tested on the left out 20% epochs and the predictions it returned were compared to the ‘true’ targets. This partitioning of the data was performed 5 times in total, each partition corresponding to a ‘fold’. The model performance in this work was averaged across folds. To estimate model performance, the sensitivity for each class was computed (corresponding to the class accuracies for ‘faces’ and ‘non-faces’). Those values were then averaged to provide a global measure of model performance, further referred to as ‘balanced accuracy’. Within each fold, another 4-folds cross-validation was performed to optimize the soft-margin hyperparameter of the SVM model (C = 0.01, 0.1, 1, 10 or 100). The significance of the obtained model performance (at p<0.05) was assessed using 1,000 permutations (58). Classification accuracy was considered significant if the balanced accuracy was significant. In addition, the difference between each model and the ‘TC’ model (I) was tested for significance based on Wilcoxon signed-rank test at the population level (n=8). To ensure a fair comparison between those models, all modeling parameters were identical, including cross-validation folds, epochs considered and permutations of the labels. Only the sets of sites considered for modeling differed across the three schemes.

### Sparse versus distributed decoding information

Assessing relevance in encoding and decoding settings assumes the interpretation of negative results. As negative results are by nature inconclusive, we here investigate the effect of priors on model performance. For one model (Model I), the algorithm used assumes a ‘distributed’ prior (SVM, L2-regularization), i.e. all features contribute to the model. If this prior is appropriate, model performance should be high, and potentially higher than other types of prior. We test this hypothesis by comparing Model I to a model that enforces sparsity on the sites. Thereby, if the sparse model performs better or as well as a distributed model, the face information is likely localized in a few sites and other features bring no further relevant information. The sparse model, further referred to as Model IV, is based on the sparse Multiple Kernel Learning method (30) described in Supplementary S2. The main difference with Model I is that sparsity is enforced at the site level. In practice, one linear kernel is built per site (i.e. number of features: 1 x 351 time points) and those kernels are combined through a sparse prior (L1-norm regularization). Some sites will hence not be considered in the final model (i.e. their contribution is zero). All modeling parameters are identical between Model I and Model IV. Furthermore, the implementation of the sparse algorithm relies on an SVM (for each kernel), which limits the effect of implementation on the results. From the output of Model IV, a ‘contribution’ map can be built, which displays the contribution of each site to the final decoding model. Interpreting contribution maps is controversial(59) as the amplitude of the contribution does not necessarily reflect the presence or absence of the signal of interest on a site. In this work, we correlate (Pearson correlation) the model contribution at each site with its human face selectivity to ensure that decoding is performed using information from face selective patches, both within and across subjects.

### Circular analysis and univariate analysis of visual localizer

In this work, we perform various analyses on the same data set, and more importantly, the contrasts investigated are identical in the univariate and multivariate analyses. The ideal solution would be to split our data set in two parts: one to identify face and human face selective sites using univariate methods, the other to build the machine learning based models. However, with numbers of trials as low as 30 for human faces after artifact rejection, splitting our data set would be detrimental for both analyses. In this section, we perform univariate analyses on another visual task recorded on the same subjects that includes human faces and non face stimuli, to investigate the amount of task dependence on the identified sites. It is important to note that this visual task comprises a different contrast from our main task, as the visual task mostly comprises images of words and numbers, and not pictures. This explains why we chose not to include this task in our main analyses.

### Experimental design

All subjects underwent a visual localizer comprising images of human faces, animal (faces with bodies), places, objects, logos, false fonts, English and Spanish words, numbers and Persian numbers. Each image was presented for 400ms, with an inter-stimulus time interval of 500ms. During the stimuli presentation, the patient was asked to pay attention to the center of the screen where a dot that would randomly change color (from red to blue) was displayed. The patient was asked to respond using a keyboard (‘press 1’) when the dot changed color.

### Analysis

This session was pre-processed similarly to the main task presented in the manuscript and univariate testing was performed to identify face selective and human face selective sites (permutation testing, FDR corrected). For face selective sites, the human face and animal categories were pooled together, while all other categories formed the non-face category. The time window considered for this analysis was amended to account for shorter presentation duration to [100 400]ms.

### Temporal distribution of face information

We implemented a technique to estimate the onset of the task-induced trial-by-trial HFB response on each site. Importantly this analysis was performed on the non-smoothed data to eliminate any confound associated with temporal smoothing. For each trial of one category, we normalize the signal with respect to peak amplitude and implement a sliding window with 30ms bins with 28ms overlap. Next, we estimate the signal average and standard deviation in a baseline time window of [-200 0]ms before onset (averaged across trials) and identify 25 consecutive bins in which the average HFB power exceeds the baseline average plus one standard deviation. This criterion allows us to identify the task-induced signal as opposed to more transient pathological activity or artefactual spiking. The earliest time point of the first bin in this sequence is marked as the signal onset for a specific trial. In the case that 25 consecutive bins surpassing the baseline threshold are not found, we exclude that trial from further analysis. In the present work, we calculate the median over trial-by-trial ROL estimates in order to assign singular ROL values for specific sites. Sites for which a ROL value could not be obtained in 50% of the trials or more were discarded from the analysis.

We investigated whether the response onset to human faces is related to the site’s anatomical position or selectivity. To this end, we computed the ROL of the HFB amplitude generated by human face stimuli on human face selective sites. The obtained ROL values (n=40) were then correlated (Spearman correlation) with the sites’ anatomical position (the ‘y’-coordinate in MNI space, estimating how posterior or anterior in the TC a site is). Similarly, the ROL values for human face selective sites were correlated (Spearman correlation) with the site’s selectivity to human faces. The same analysis was performed for task active sites (n=94) for comparison. Our ROL analyses are subject to imprecisions, due to temporal smoothing related to the time-frequency decomposition and to the averaging of the HFB signal in 30ms bins. Hence our focus is on estimating relationships between ROL and anatomical positions. Absolute values of ROL should be considered with care as different techniques lead to different ROL values (60). To this end, we estimated the same correlations but subtracting the ROL value from the most face selective site within each subject. These results, plotted in Figure 4b,c and S4a,b ensure that the reported correlations are not driven by specific subjects or systematic errors in timing estimation.

### Effects of signal amplitude and slope on ROL

Our method performs response onset detection at the trial level, based on unsmoothed data. However, different parameters of the signal could affect the obtained ROL values, including noise, signal amplitude and signal slope. In this section, we used semi-simulated data to investigate the effect of signal amplitude and slope on the obtained ROL values. The level of noise is the one that is naturally present in ECoG data.

The semi-simulated data used in this work have been designed for other work(61) and are described in detail below. The data and code for generating the simulation are available open-source (https://github.com/JessicaSchrouff/Simulated_ECoG).

### Original data

The data was recorded from Subject S1, during a 5-minute wakefulness rest period, with eyes closed. Sites assessed as ‘pathological’ by medical doctors were discarded from further analysis.

### Simulated design

A fake experimental design was simulated: 2 conditions, ‘A’ and ‘B’, presented at random every 1.9 seconds. The stimuli are further assumed to last for 1 second. This yielded 146 stimuli, 73 for each category.

### Preprocessing

Signal pre-processing was performed with specific ECoG routines (github/LBCN/Preprocessing_Pipeline) using Matlab (www.mathworks.com) and SPM12 (www.fil.ion.ucl.ac.uk/spm). First, the data was converted to SPM format and downsampled to 1kHz. The continuous signal was filtered for line noise and harmonics (stop-band: 57-63Hz, 117-123Hz, 177-183Hz) and an automatic quality assessment identified ‘noisy’ or ‘spiky’ sites based on their variance and number of ‘jumps’ (i.e. signal derivative>100µV), leaving 38 ‘good’ sites. The data was re-referenced to the average of all good channels before being epoched in the [-400, 1400]ms window around ‘onset’ and baseline corrected using the [-400, 0]ms window. Epochs displaying flat segments of more than 4ms or ‘jumps’ larger than 100µV were discarded from further analysis. The signal was then decomposed using a 5-wavelets decomposition in the 70 to 170Hz frequency band (step: 10Hz, avoiding 120Hz) to estimate High Frequency Broadband (HFB) power. The time-frequency signal was z-scored based on the pooled baselines of all events in the [-300, 0]ms window before onset to avoid edge effects and smoothed in the [-200, 1200]ms window after onset by a 50ms Gaussian window. Epochs displaying z-scores larger than 8 were discarded, leaving 60 trials for condition ‘A’ and 56 for ‘B’. This pre-processing procedure is very similar to the one used in the main part of the work, including bad channel rejection.

### Simulated signals

All modifications of data structure were performed on the pre-processed data to avoid an effect of the pre-processing on the obtained results. To simulate neural signal, a ramp window was added to all epochs of condition ‘A’ starting 0ms after ‘onset’ with a slope of 3 until 500ms, on all ‘good’ sites. The amplitude of the signal in condition ‘A’ was varied by modifying the Signal-to-Noise Ratio (SNR) between trials ‘A’ and ‘B’. Hence, varying signal amplitude is strongly correlated with varying the selectivity of sites to condition ‘A’. The amplitude of the signal in the ramp window was computed based on a desired SNR on each site:

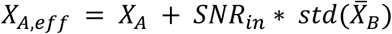

Where *X*_*A,eff*_represents the amplitude of the effective simulated signal for condition ‘A’ trials, *X*_*A*_, the amplitude of the signal for trials ‘A’, *SNR*_*in*_, a fixed number representing the desired SNR and 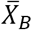, the average trace of B trials. *SNR*_*in*_ was varied from 2 to 10 by steps of 0.5. In our real dataset, the distribution of estimated SNR varies from −2 to 17 on human face selective sites, with only 3 sites with SNR>10.

To estimate the effect of signal slope on the ROL results, we performed the same simulation but normalizing the amplitude of the signal in each trial before ROL detection (Linf-norm, i.e. dividing by the maximum amplitude). This simulated dataset hence varies the slope of the signal, but not the amplitude (set to maximum of 1).

### ROL analysis

For each SNR level, ROL detection is performed at the trial level, for the ‘A’ trials on each ‘good’ site. All parameters are identical to the technique reported above. Trials with less than 50% of detected onsets were excluded from the results. This analysis was performed on both the un-normalized (i.e. varying amplitude at fixed slope) and the normalized (i.e. varying slope at fixed amplitude) simulated data.

### Causal importance of face selective sites for face perception

A set of electrical brain stimulations (EBS) was performed on seven of the eight subjects. The sites of stimulations were chosen based on a priori knowledge about their HFB responses to face and non-face stimuli. During the procedure, face selective and task active sites were stimulated with electrical charge while the subjects were instructed to look at various real-world face stimuli (persons at the bedside). Across the seven subjects, the instructions given included A) looking at the face, B) looking at the lips, C) looking at the nose, D) looking at self in the mirror, E) looking at a cartoon face on a sheet of paper, or F) close eyes and imagine a face. Between 3 to 8 mA (depending on the excitability of the stimulated site) were delivered at a duration ranging from 1 to 3 seconds at 50 Hz frequency and 200 μs pulse width, of a square wave electrical waveform in unipolar (subject S8) or bipolar montage (S8 and other subjects). In unipolar montage a TC site and a remote cortical reference site were stimulated whereas in the bipolar montage, a pair of adjacent electrodes were stimulated. Sham stimulations were also administered at 0 mA. Continuous EEG monitoring showed no after-discharges or epileptic activity during the sessions. Verbal reports were collected following each stimulation. See Figure 5 legend for criteria used to define a site as active during EBS procedure. We recognized an electrode as a catalyst in face perception change if A) the stimulation of this electrode yielded a face-specific change; B) the stimulation of this electrode paired with any other electrode still yielded a change; and C) the result of the stimulation and its resulting face-perception change was replicable across multiple stimulation trials. In addition, in subject S8, the estimated cortical area affected by the stimulation on electrode 3 was calculated using the relationship between the estimated charge per trial and the cortical area affected(62) and is shown with a blue circle around it. The charge deposited per trial (μC) was calculated as a product of the pulse width (ms), current (mA), frequency (Hz), and duration (s) of stimulation for each trial; then, we estimated the cortical area (mm^2^) affected by the stimulation as a function of the charge deposited per trial (μC) according to the methods described in (62).

### Statistical testing

Throughout this work, statistical testing was performed using non-parametric permutations. When suited, the tests were paired (e.g. when comparing 2 conditions in terms of ROL value on the same set of sites, or when comparing the HFB response of a stimulus category to its baseline). A minimum of 1,000 permutations was performed, to ensure a good estimation of the null distribution. False Discovery Rate (FDR) or Bonferroni correction was applied when multiple comparisons tested for the same effect (e.g. testing for human face selectivity on each site, or comparing ROL between the 4 face subcategories using binary comparisons). Significance was determined at p<0.05, after correction if applicable. Population statistics for the decoding models I to IV was performed using Wilcoxon signed-rank tests (58).

## SUPPLEMENTARY INFORMATION

## Supplementary figures

**Figure S1:**
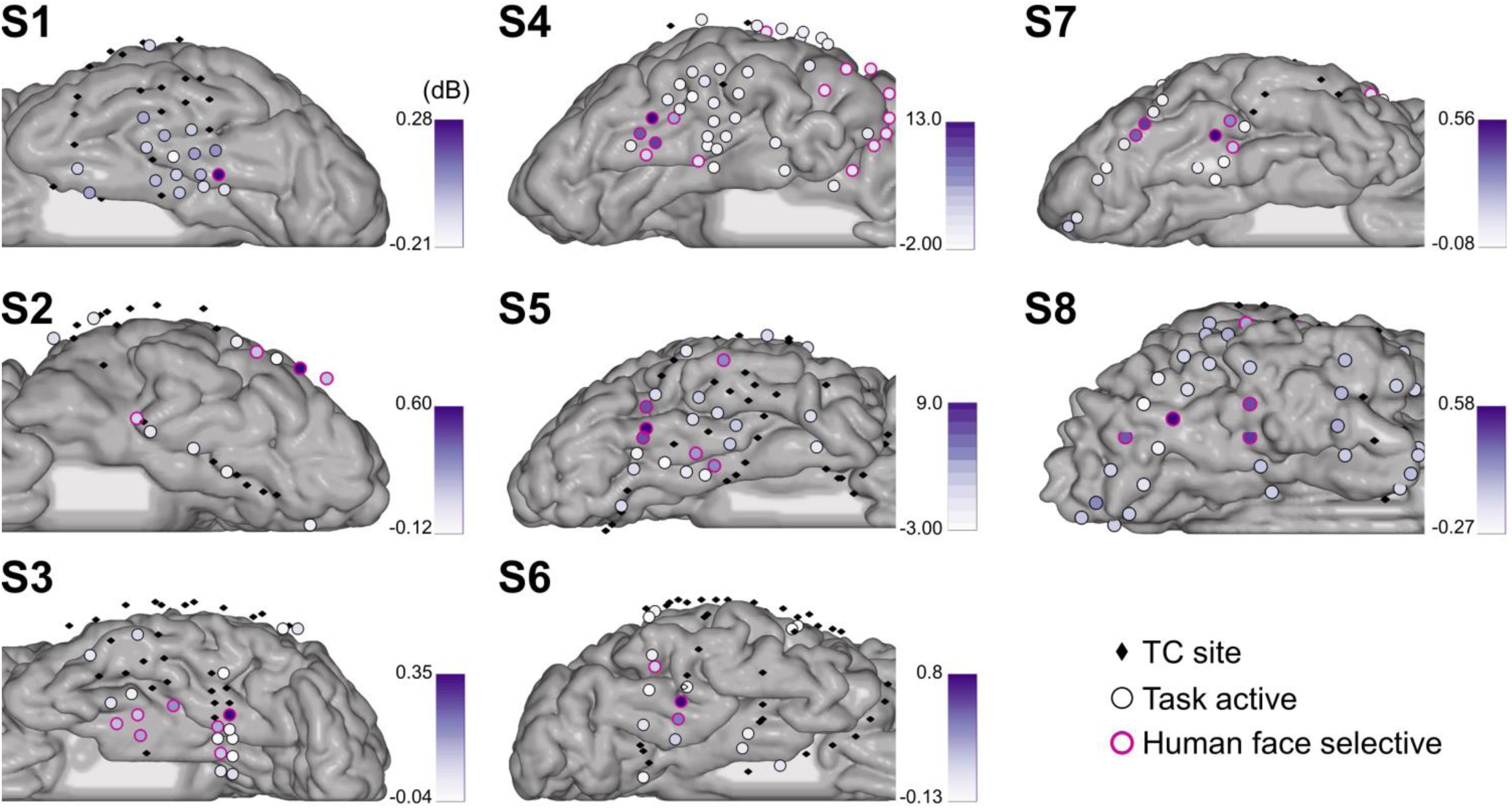
Human face selective sites, within subjects. For each subject, the sites are represented on the individual cortex map. Sites represented by black diamonds were not assessed as task active. The first three subjects have left hemisphere implantation, while subjects S4 to S8 have right implantation. Sites displayed by circles were assessed as task active. Among those, sites highlighted by a pink rim were further assessed as human face selective. The human face selectivity of each site (in dB) is displayed via a color-coded fill of the task active sites.

**Figure S2:**
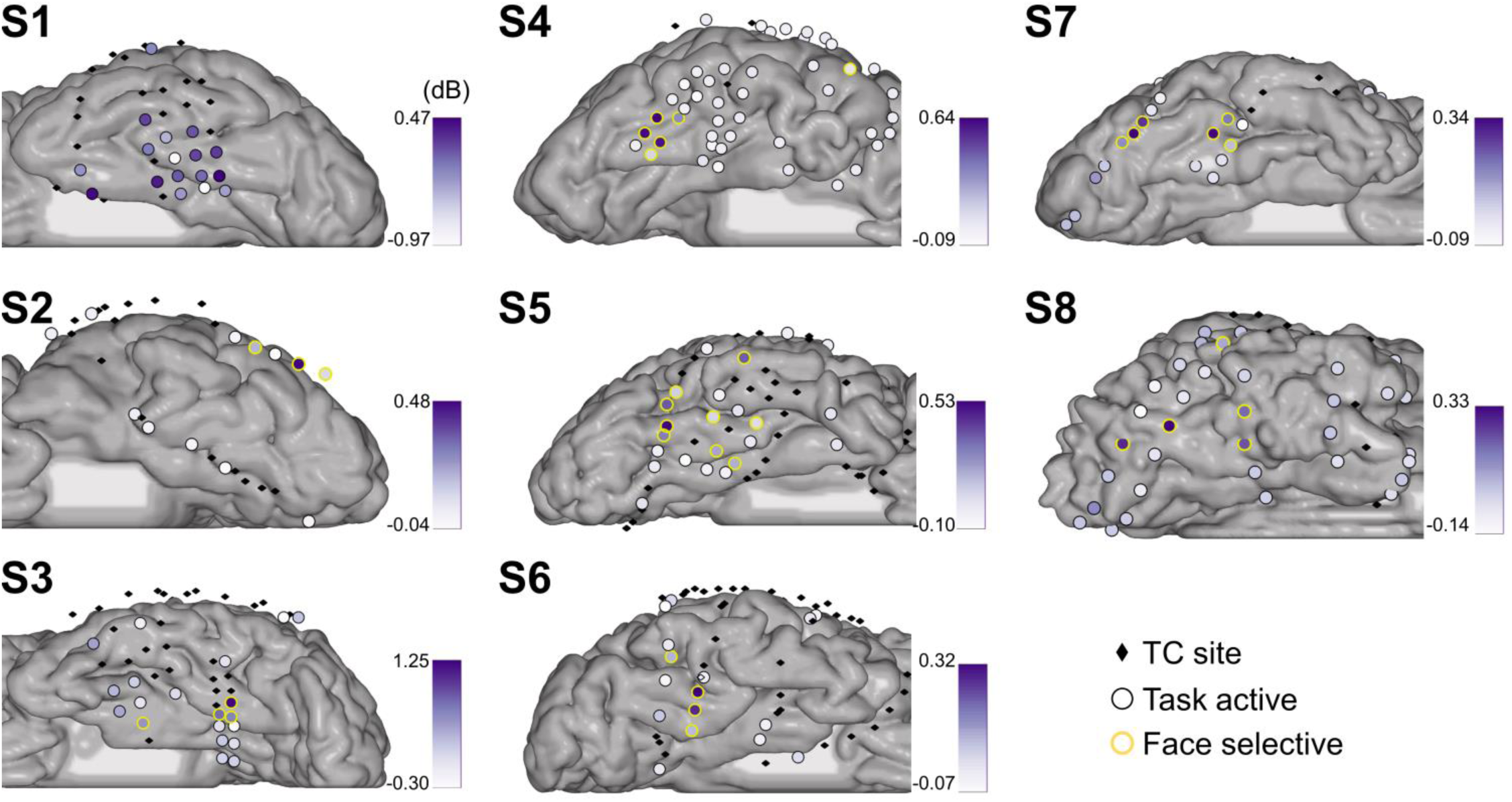
Face selective sites, within subjects. For each subject, the sites are represented on the individual cortex map. Sites represented by black diamonds were not assessed as task active. The first three subjects have left hemisphere implantation, while subjects S4 to S8 have right implantation. Sites displayed by circles were assessed as task active. Among those, sites highlighted by a yellow rim were further assessed as face selective (four face subcategories pooled). The face selectivity of each site (in dB) is displayed via a color-coded fill of the task active sites.

**Figure S3:**
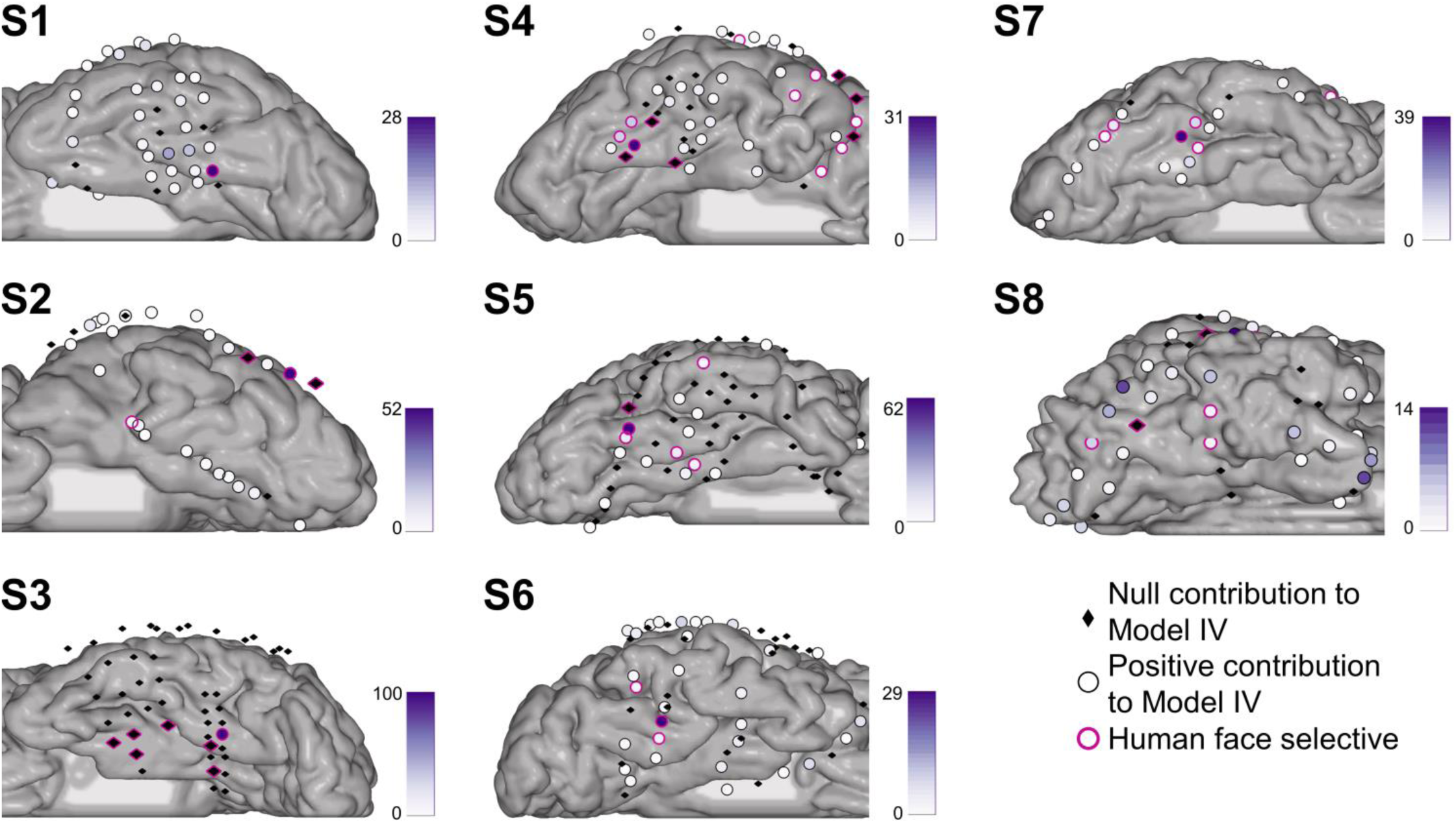
Individual plots of site contribution to Model IV. For each subject, the sites are displayed on the individual cortex mesh. Sites represented by black diamonds have a null contribution to Model IV. Sites represented by a circle have a positive contribution to Model IV, the amplitude of their contribution being color-coded using a purple fill. Sites assessed as human face selective by our univariate analysis are highlighted with a pink rim.

**Figure S4:**
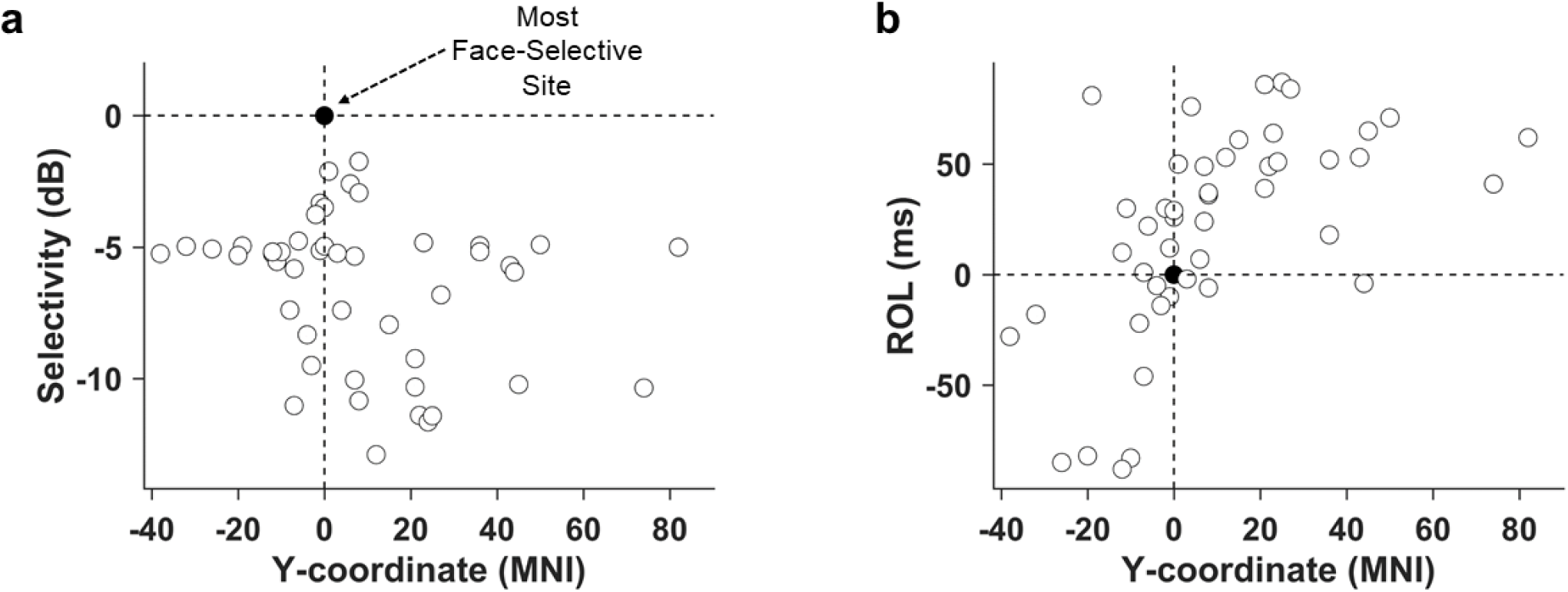
Temporal distribution of human face information on task active sites. **(a)** Selectivity in each subject, when compared to MNI y-coordinate with the best (i.e. most human face selective) site in each individual chosen as the point of reference. The best site is represented as a black circle, at the crossing of the 2 axes. Selectivity is not related to anatomical position. **(b)** ROL in each subject, when compared to MNI y-coordinate with the best site in each individual chosen as the point of reference. Latency is related to anatomical location of the electrodes.

**Figure S5:**
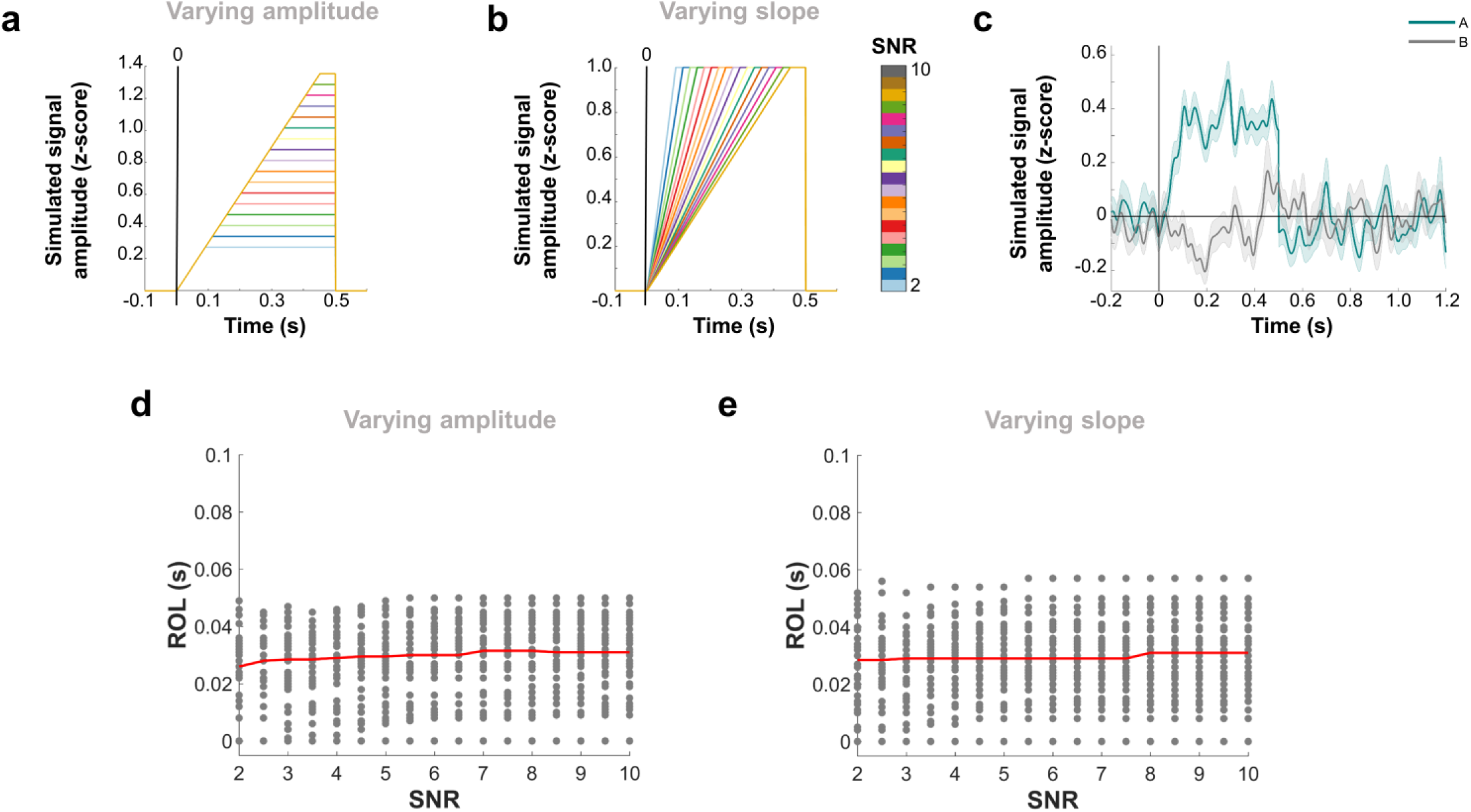
Effect of signal amplitude and of slope on ROL. **(a)** Simulated ramp signals, for varying imposed SNR, for an example site. The onset time is 0ms after ‘onset’. In the present case, the slope is fixed and the amplitude varies with imposed SNR. **(b)** Simulated ramp signals after normalizing by the maximum amplitude (i.e. Linf-norm), for the same example site. In this case, the slope varies but the maximum amplitude is fixed at 1. **(c)** Example obtained semi-simulated signals for one site, at imposed SNR = 3. The average trace for condition ‘A’ is displayed in green (with shaded standard error), and condition ‘B’ (unused in this work), in grey. The amount of noise represented is the on-going resting activity from the recorded data. **(d)** When varying amplitude at fixed slope (i.e. un-normalized signals), SNR does not affect significantly the detected ROL across sites (ρ = 0.0613, p = 0.1198). **(e)** Similar results are obtained when varying slope at fixed amplitude, i.e. normalizing the signals (ρ = 0.0245, p = 0.5335). Please note that the effect of noise is displayed by the variability in ROL across sites for each SNR.

### Supplemental Information

**Supplementary Table S1:** Demographics. *** Because we are enclosing a video of this subject, we are keeping his information confidential.

| Subject # | Age | Gender | Native Language | Language Lateralization | Handedness | Type of epilepsy | Side of implantation (Number of electrodes) | IQ | Duration of epilepsy |
| --- | --- | --- | --- | --- | --- | --- | --- | --- | --- |
| 1 | 47 | male | English | Bilateral | left | Right temporal lobe epilepsy | left (64) | 74 | 13 years |
| 2 | 44 | male | Spanish | N/A | right | Left Temporal lobe epilepsy | left (106) | N/A | 41 years |
| 3 | 23 | male | English | Left | right | Left temporal oligodendroglioma | left (125) | 100 | 6 years |
| 4 | 68 | male | Korean | N/A | right | Right temporal lobe | right (96) | N/A | 32 years |
| 5 | 65 | female | English | Right | right | Right temporal lobe epilepsy | right (110) | 113 | 4 years |
| 6 | 35 | male | German; fluent in English | Left | right | Right fronto-temporal lobe | right (126) | 129 | 18 years |
| 7 | 36 | female | English | N/A | right | Right temporal lobe epilepsy | right (128) | 102 | 6 years |
| 8 | *** | male | English | *** |  |  |  |  |  |

### Supplementary 2: Frequency information for faces

Table S2: suggests that other frequency bands carry (human) face information. However, the MKL model strongly prefers the HFB to discriminate human faces from non-faces, in all subjects (Figure 2b). This result suggests that either the human face information carried by other bands is weaker, or it is highly correlated with the HFB human face information (as the MKL model only selects non-correlated information). Therefore, using the amplitude of the HFB power as an index of neural population activity for investigating (human) face distribution over the TC seems reasonable.

**Supplementary table S2:** Task active and (human) face selective sites in each frequency band. The number of identified task active, face selective and human face selective sites is displayed, summed over all subjects.

| Band | Task active | Face selective | Human Face selective |
| --- | --- | --- | --- |
| $\theta$ | 122 | 2 | 2 |
| $\alpha$ | 104 | 5 | 7 |
| $\beta_1$ | 98 | 17 | 25 |
| $\beta_2$ | 80 | 9 | 23 |
| $\gamma$ | 114 | 21 | 30 |
| HFB | 193 | 37 | 48 |

**Supplementary Table S3:** TC sites and category-specific responses. For each participant, the number of sites located in the TC is displayed. Sites were excluded from further analysis if they exhibited epileptic activity or excessive noise artifacts. The number of task active (3^rd^ column), face selective (4^th^ column) and human face selective (last column) sites are also displayed. The total for each column is displayed at the bottom of the table.

| Patient | TC sites | Task active | Face selective | Human face selective |
| --- | --- | --- | --- | --- |
| S1 | 39 | 17 | 0 | 1 |
| S2 | 29 | 12 | 3 | 4 |
| S3 | 50 | 20 | 4 | 7 |
| S4 | 49 | 45 | 6 | 15 |
| S5 | 59 | 23 | 9 | 6 |
| S6 | 55 | 16 | 4 | 3 |
| S7 | 30 | 21 | 6 | 6 |
| S8 | 46 | 36 | 5 | 6 |
| **Total** | **357** | **190** | **37** | **48** |

### Supplementary S4: Low-level image features

#### Stimulus features

There were no significant differences in spatial information when comparing the 11 categories, pair-wise (p-values = 0.9999). Mean luminance was significantly different between human faces and non-faces (p=9.93e-09, n= 200), mammal faces and non-faces (p=6.3e-08, n=200), bird faces and non-faces (p=4.5e-04, n=200) and marine faces and non-faces (p=6.2e-07, n=200). No significant differences were found between human and mammal faces (p=0.88, n= 25), human and bird faces (p=0.15, n=25), human and marine faces (p=0.71, n=25), mammal and bird faces (p=0.67, n=25), mammal and marine faces (p=0.99, n=25) and bird and marine faces (p=0.84, n=25).

**Supplementary Table S4:**
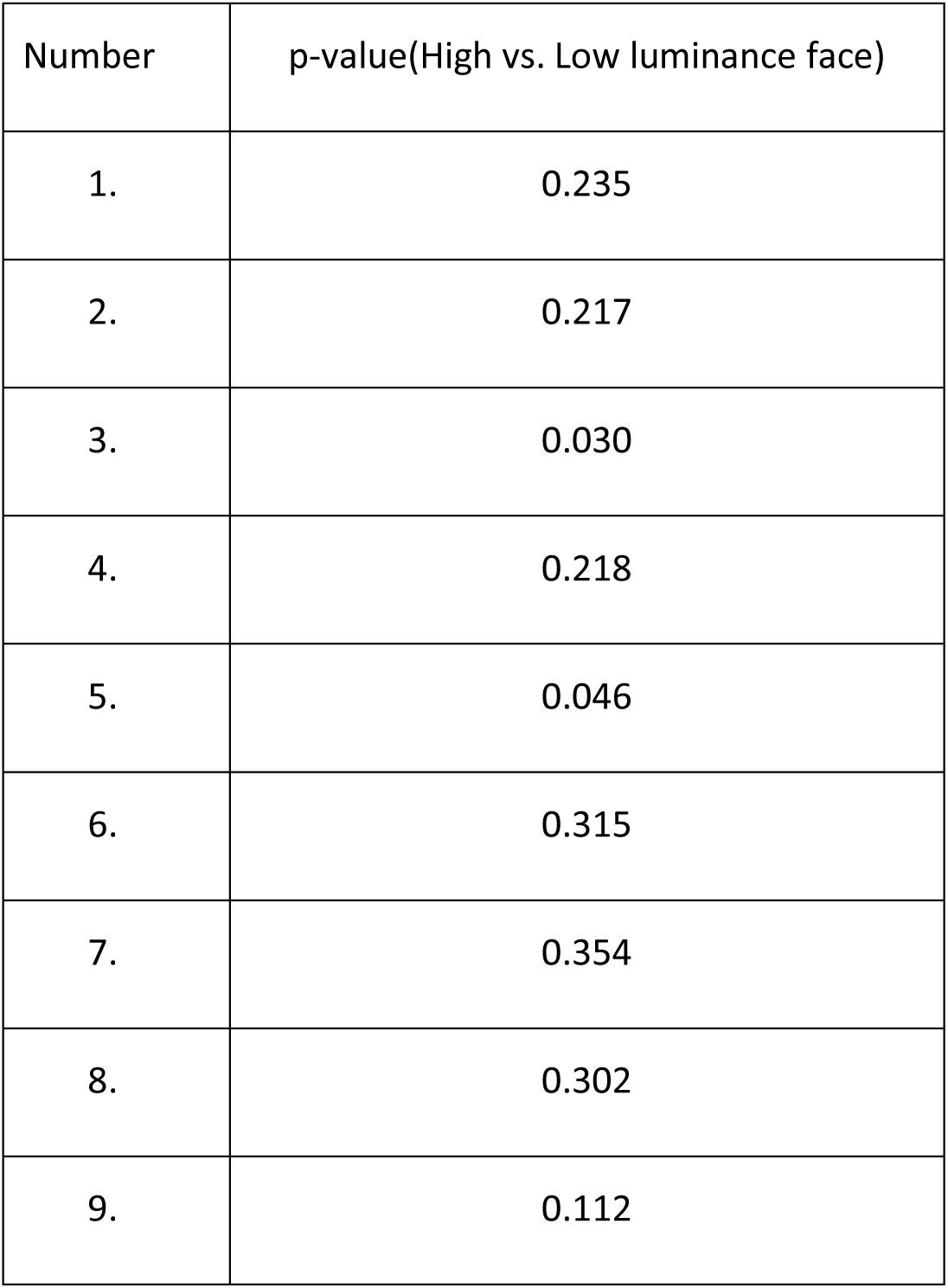
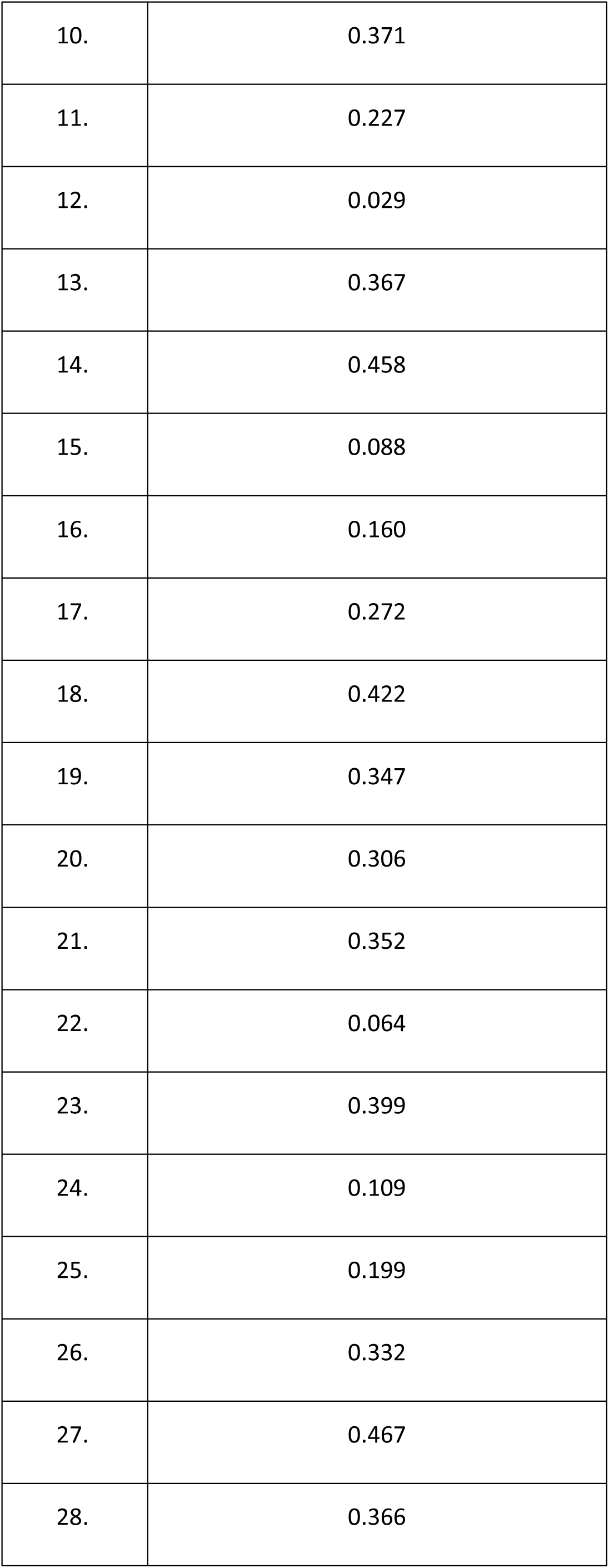
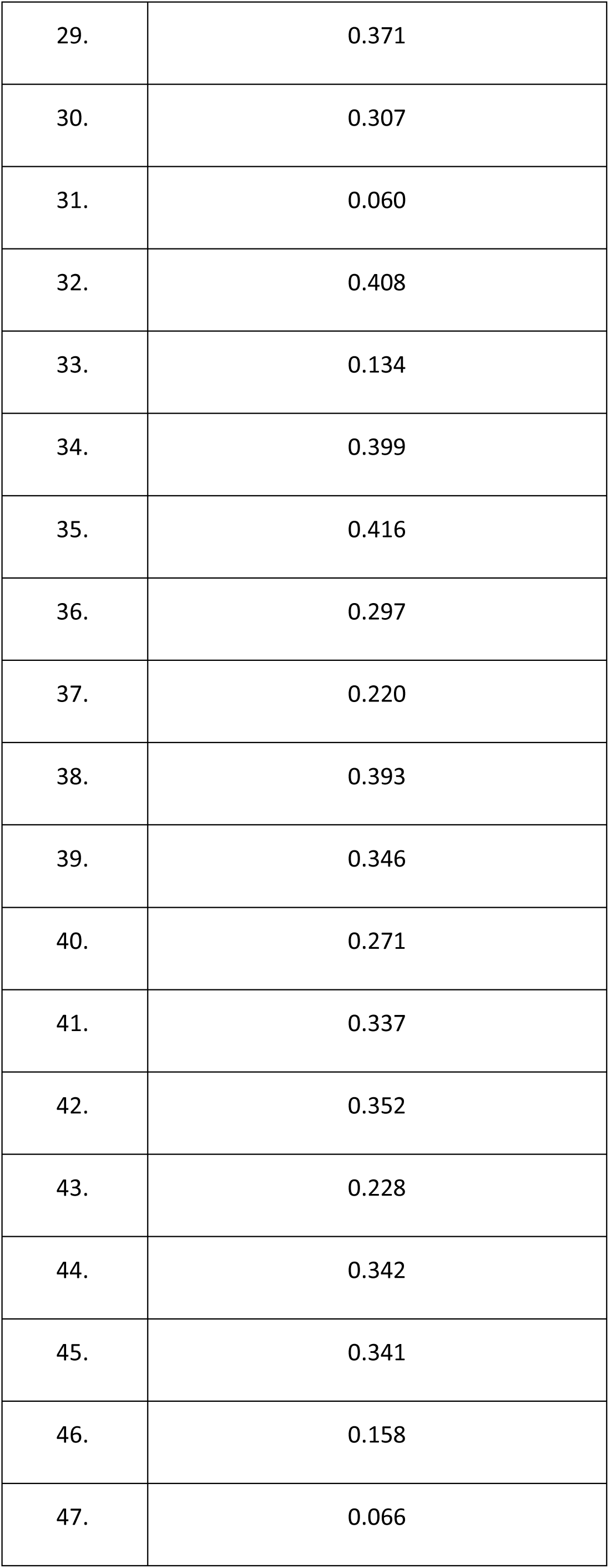
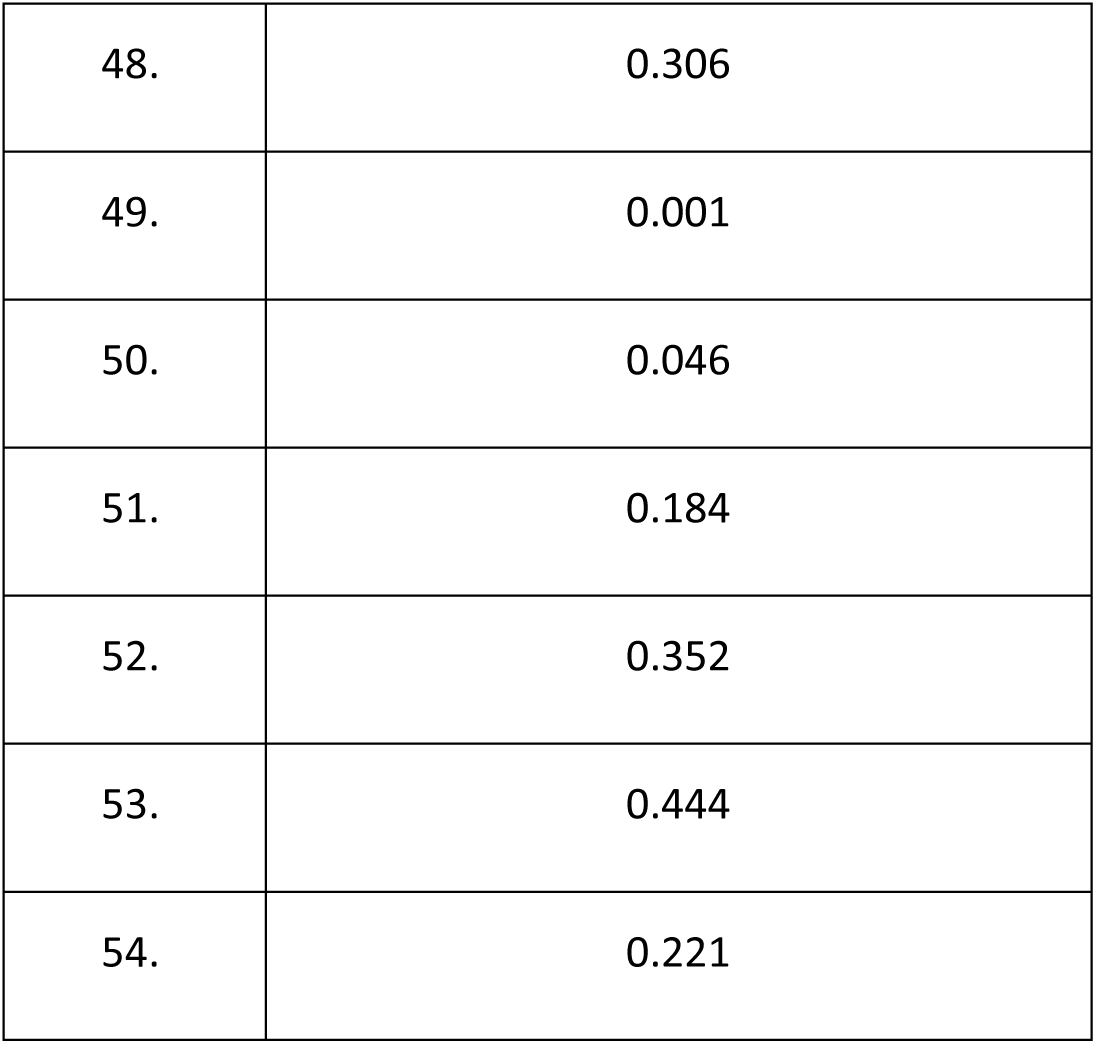
Effect of luminance on face sites. P-values of univariate analyses comparing low luminance face stimuli with high luminance face stimuli.

| Number | p-value(High vs. Low luminance face) |
| --- | --- |
| 1. | 0.235 |
| 2. | 0.217 |
| 3. | 0.030 |
| 4. | 0.218 |
| 5. | 0.046 |
| 6. | 0.315 |
| 7. | 0.354 |
| 8. | 0.302 |
| 9. | 0.112 |
| 10. | 0.371 |
| 11. | 0.227 |
| 12. | 0.029 |
| 13. | 0.367 |
| 14. | 0.458 |
| 15. | 0.088 |
| 16. | 0.160 |
| 17. | 0.272 |
| 18. | 0.422 |
| 19. | 0.347 |
| 20. | 0.306 |
| 21. | 0.352 |
| 22. | 0.064 |
| 23. | 0.399 |
| 24. | 0.109 |
| 25. | 0.199 |
| 26. | 0.332 |
| 27. | 0.467 |
| 28. | 0.366 |
| 29. | 0.371 |
| 30. | 0.307 |
| 31. | 0.060 |
| 32. | 0.408 |
| 33. | 0.134 |
| 34. | 0.399 |
| 35. | 0.416 |
| 36. | 0.297 |
| 37. | 0.220 |
| 38. | 0.393 |
| 39. | 0.346 |
| 40. | 0.271 |
| 41. | 0.337 |
| 42. | 0.352 |
| 43. | 0.228 |
| 44. | 0.342 |
| 45. | 0.341 |
| 46. | 0.158 |
| 47. | 0.066 |
| 48. | 0.306 |
| 49. | 0.001 |
| 50. | 0.046 |
| 51. | 0.184 |
| 52. | 0.352 |
| 53. | 0.444 |
| 54. | 0.221 |

**Supplementary Table S5:** Relevance of face and non-face sites in decoding. The first column displays the balanced accuracy for Model I, with the p-value that it differs from the distribution defined by the ‘random set’ models. The second column displays the performance of the same model when discarding human face selective sites (Model II), with the p-value that it differs from the ‘random set’ models. Column 3 displays the balanced accuracy of the same model when random subsets of sites are discarded (‘random set’, median across 499 models). Finally, column 4 displays the correlation between the accuracy of ‘random set’ models and the proportion of human face sites included in the model. Significant results are displayed in bold (p<0.05, corrected). Results displayed in italic show a trend (p<0.05 uncorrected).

| Patient | ‘TC’ (I, in %) | ‘TC-sign’ (II, in %) | ‘random set’ (in %) | Corr(F,acc) |
| --- | --- | --- | --- | --- |
| S1 | **68.00** ($p=0.4980$) | 61.71 (**$p=0.0020$**) | 68.00 | N.a. |
| S2 | **85.54** ($p=0.3000$) | 62.82 ($p=0.0680$) | 84.64 | **0.4697** |
| S3 | **75.22** ($p=0.2340$) | 58.27 (**$p=0.0060$**) | 72.52 | **0.2441** |
| S4 | **94.98** ($p=0.7620$) | **69.48** (**$p=0.0020$**) | 95.58 | **0.2060** |
| **S5** | **98.08** (p=0.1340) | 53.89 (**p=0.0020**) | 98.00 | **-0.2144** |
| **S6** | **80.10** (p=0.6180) | **73.41**(p=0.0020) | 80.55 | *0.0971* |
| **S7** | **85.86** (p=0.6080) | 64.16 (**p=0.0020**) | 86.82 | **0.4119** |
| **S8** | **94.10** (p=0.0580) | 60.45 (**p=0.0020**) | 91.59 | **0.2611** |

#### Univariate analysis

When comparing low and high luminance faces, no site displayed a significant difference between the two categories (5 before FDR correction, Table S4). These results need to be considered with care, as these negative results cannot completely reject the null hypothesis (they might be under-powered). However, the obtained results suggest that there is no effect of luminance on our main findings.

### Supplementary S6: Univariate results on visual localizer

Table S6 displays the number of sites identified as face and human face selective on this task, as well as the number of sites commonly identified in both tasks. It reveals that 60.53% of sites identified on the main task were reported as face selective in both tasks, while 58.33% of sites were reported as human face selective in both tasks. Although the two tasks have fundamentally different contrasts (the visual localizer contains more images of words than of pictures, and the animal category contains both face and body), the overlap of selected face and human face sites is significant. It is therefore unlikely that our reported results are driven by task specific features.

**Supplementary table S6:** **Consistence of face and human face selective sites identification across tasks.** Number of face and human face selective sites identified on a visual localizer task (‘Loc’), and identified on both our main task and the visual localizer (‘both’).

| Patient | Face selective (Loc) | Face selective (both) | Human face selective (Loc) | Human face selective (Both) |
| --- | --- | --- | --- | --- |
| S1 | 6 | 0 | 3 | 1 |
| S2 | 3 | 2 | 2 | 2 |
| S3 | 5 | 2 | 2 | 2 |
| S4 | 2 | 0 | 8 | 6 |
| S5 | 9 | 5 | 16 | 5 |
| S6 | 10 | 4 | 6 | 3 |
| S7 | 10 | 7 | 7 | 5 |
| S8 | 7 | 3 | 5 | 4 |
| **Total** | **52** | **23** | **49** | **28** |

**Supplementary table S7:** Distributed versus sparse anatomical prior. The first column copies the accuracy from Model I (balanced accuracy, in %), for easy comparison. The second column displays the balanced accuracy (in %) obtained from a sparse Multiple Kernel Learning model. The third column displays the correlation (with p-value) between site contributions to the model and the human face selectivity as identified by the univariate test ‘human faces versus pooled non-faces’.

| **Patient** | **‘TC’ (I, in %)** | **sMKL (IV, in %)** | **Corr(beta,uni)** |
| --- | --- | --- | --- |
| **S1** | **68.00** | **76.95** | **0.4426** (p=0.0048) |
| **S2** | **85.54** | **90.71** | **0.6714** (p<0.0001) |
| **S3** | **75.22** | **80.07** | **0.6773** (p<0.0001) |
| **S4** | **94.98** | **96.00** | **0.7635** (p<0.0001) |
| **S5** | **98.08** | **96.15** | **0.8681** (p<0.0001) |
| **S6** | **80.10** | **77.57** | **0.7272** (p<0.0001) |
| **S7** | **85.86** | **90.86** | **0.6855** (p<0.0001) |
| **S8** | **94.10** | **91.78** | -0.1405 (p=0.3517) |

**Supplementary table S8:** Effects of signal amplitude and slope on ROL Examples of simulated ramp signals and obtained semi-simulated signals are presented in figure S5 (a-c), on one example site. Figure 5a displays the simulated signal obtained for different SNRs when varying amplitude while Figure 5b displays the simulated signal obtained for different SNRs when varying slope (i.e. normalizing the signal). The ROL-SNR relationships are displayed in Figure S4 for the un-normalized (d) and normalized signals (e). When varying amplitude at fixed slope, correlation between detected onset and estimated SNR was close to 0 (ρ = 0.0613, p = 0.1198) across all 38 sites (displayed as grey dots on the scatter plot). When varying slope at fixed amplitude, similar results were found (ρ = 0.0245, p = 0.5335). In either case, the average ROL varies at most by 6 (un-normalized) or 12 (normalized) ms between SNR = 2 and SNR =10. This scale of variation is much smaller than the effect reported between posterior and anterior sites in the main text. The proposed ROL detection method is independent of signal amplitude and slope. In addition, the size of a potential bias is limited and much inferior to the effect reported in the main text. In conclusion, it is unlikely that the reported effects result only from biases in our ROL method.

**Supplementary table S9:**
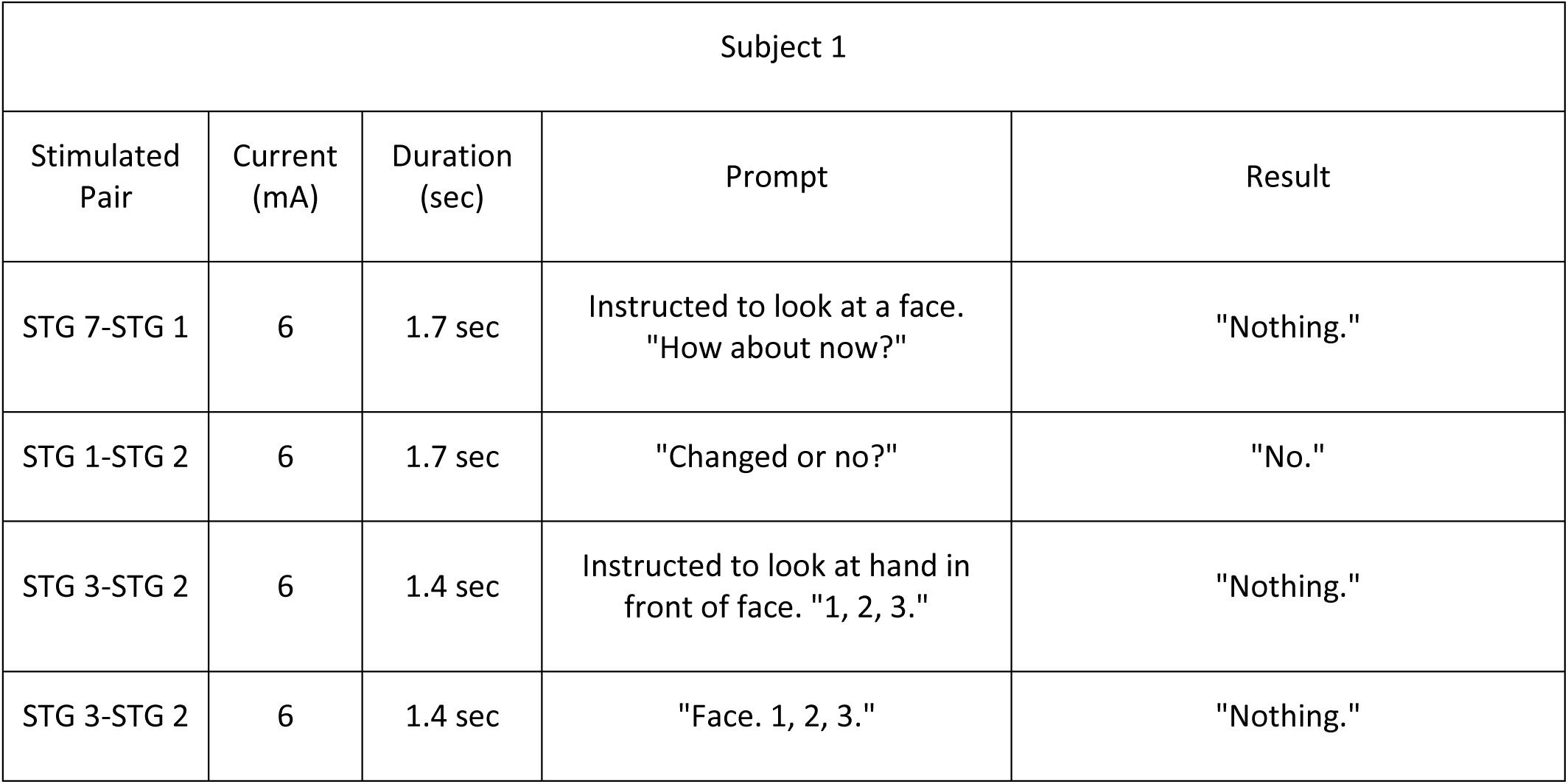
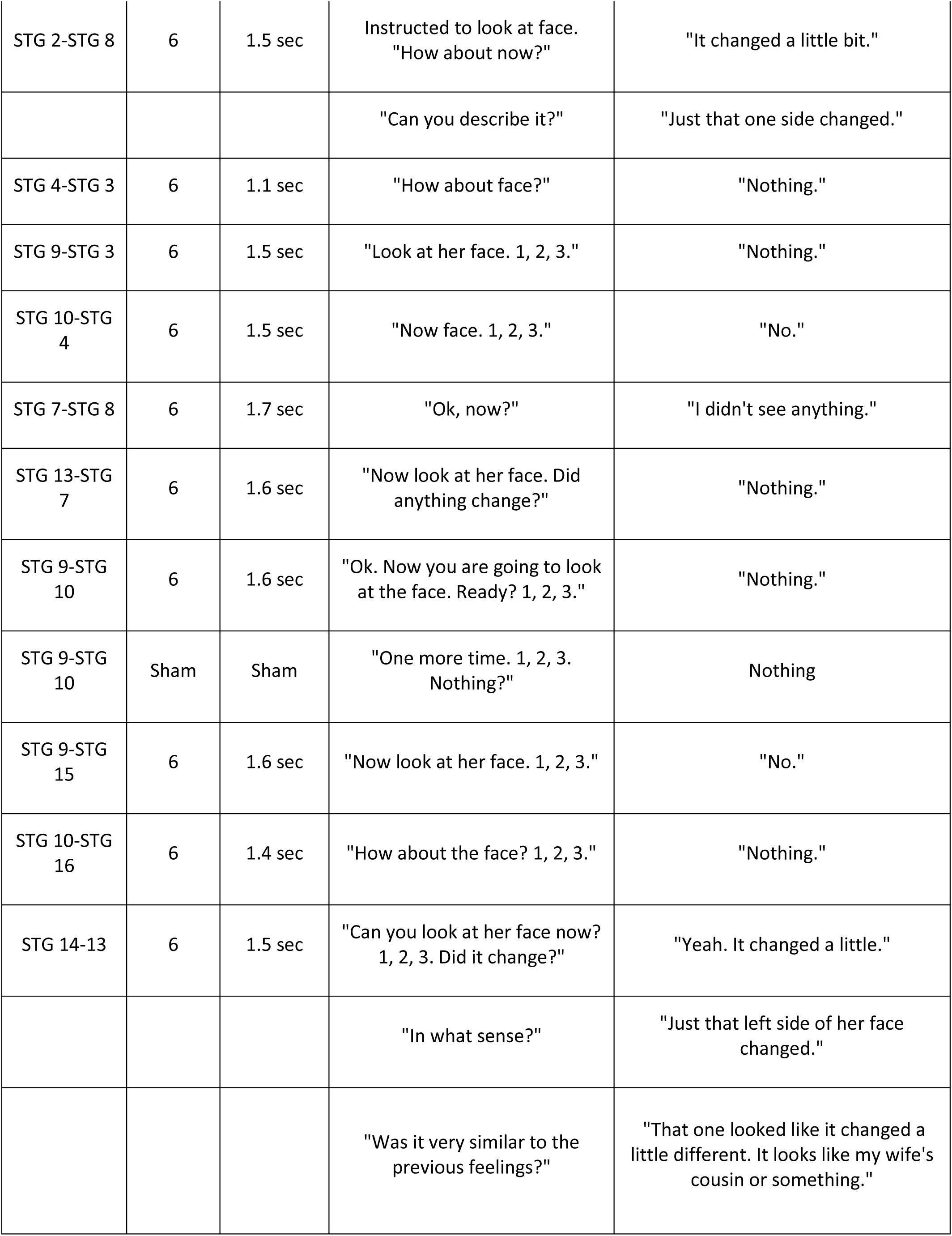
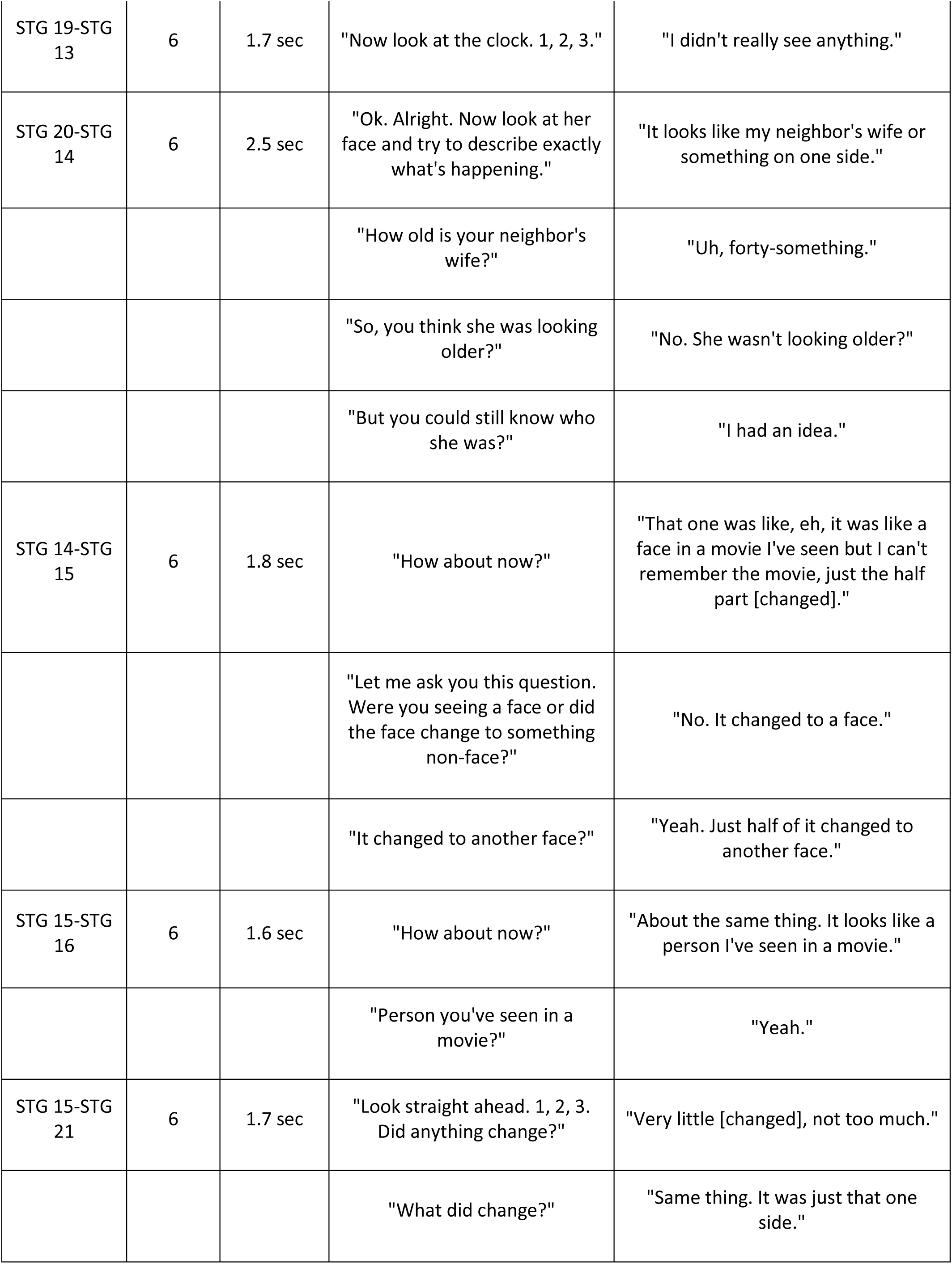
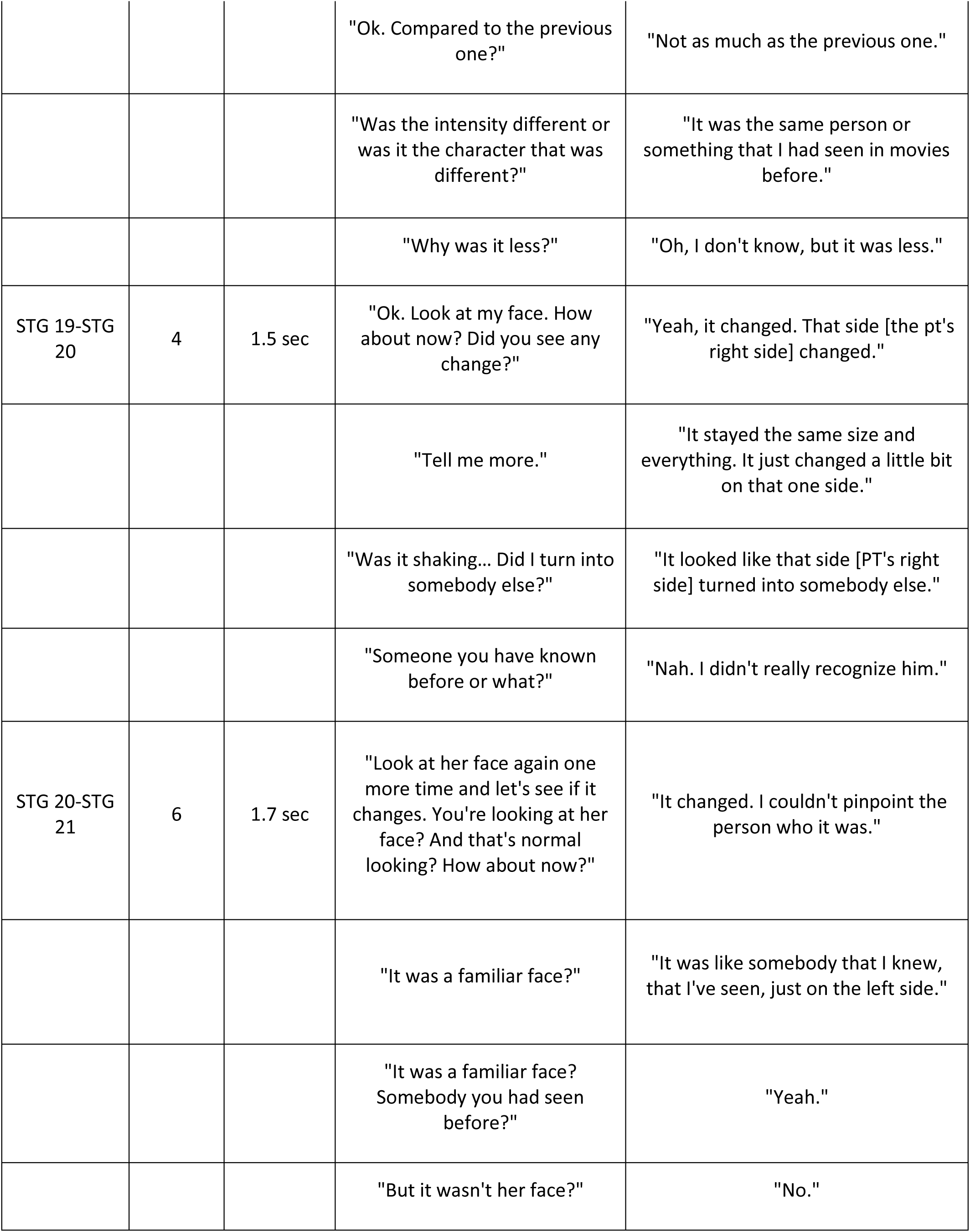
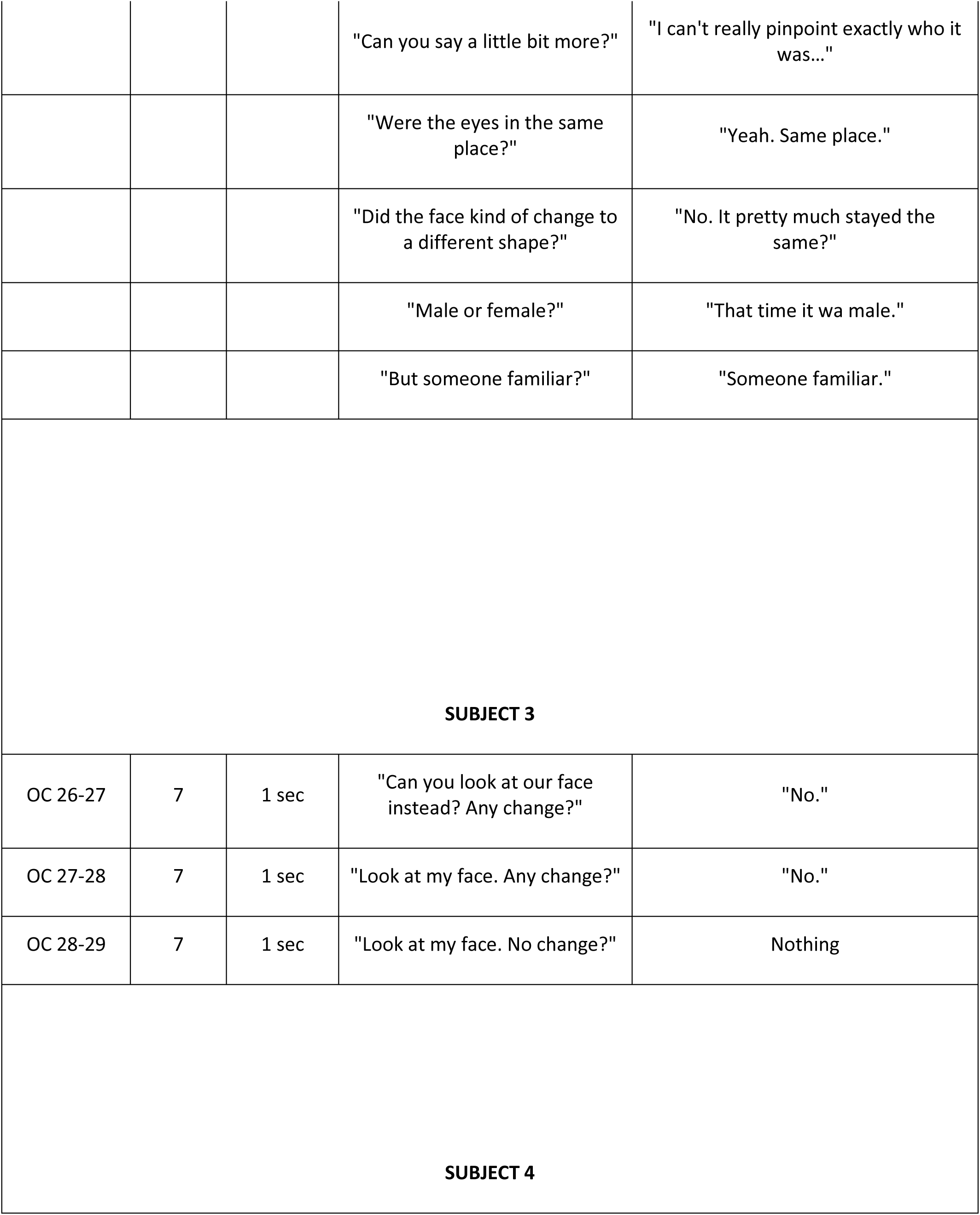
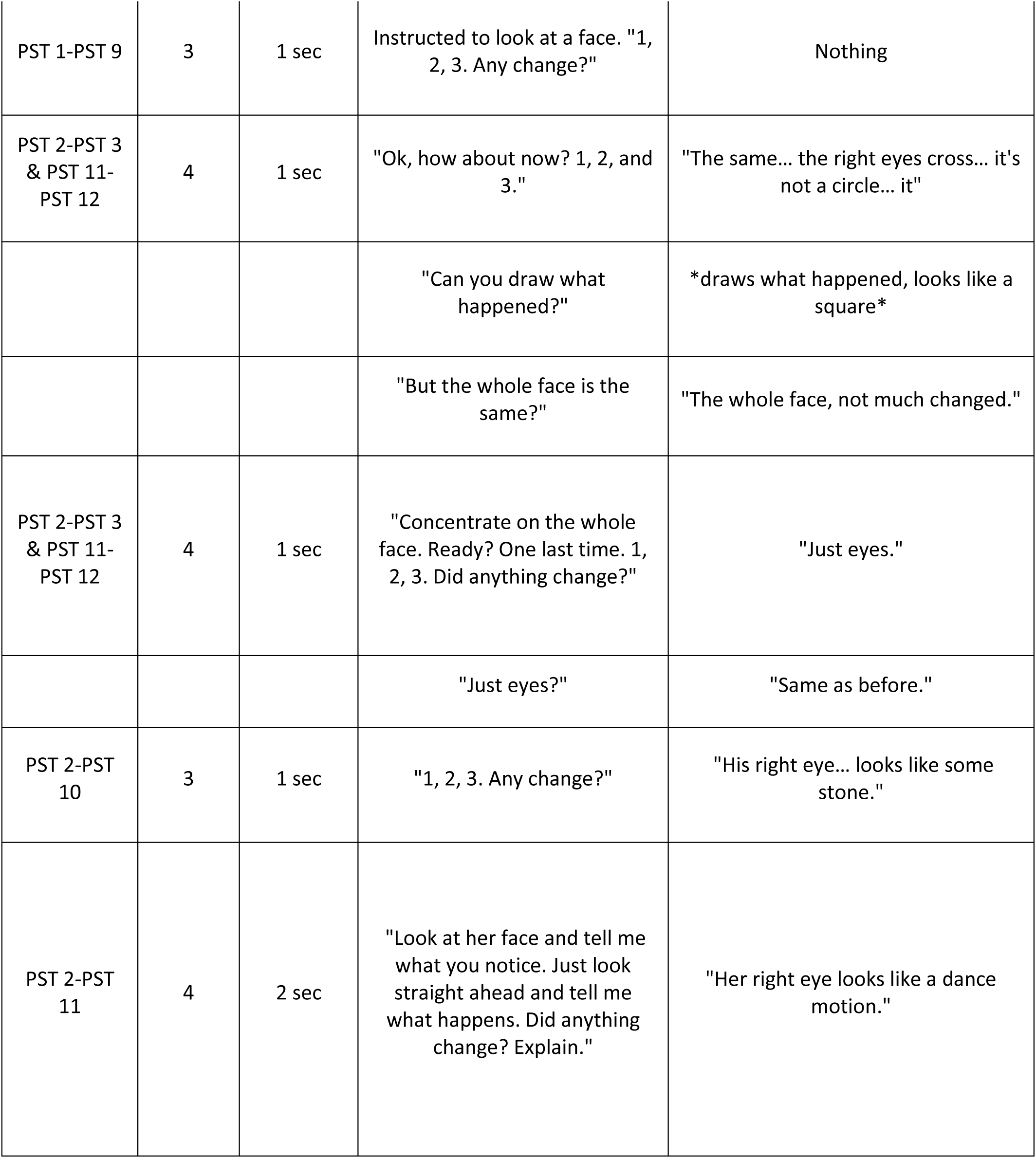
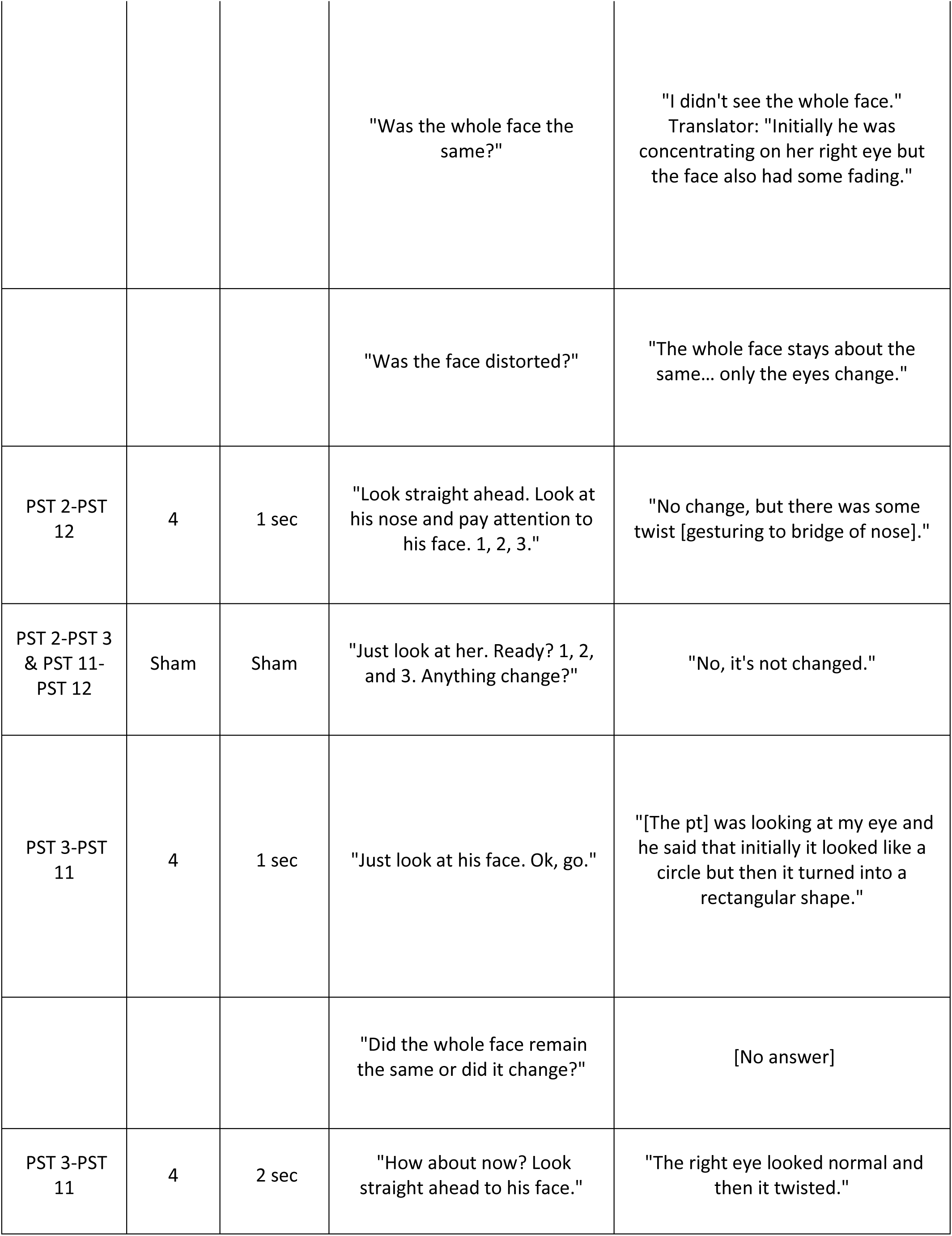
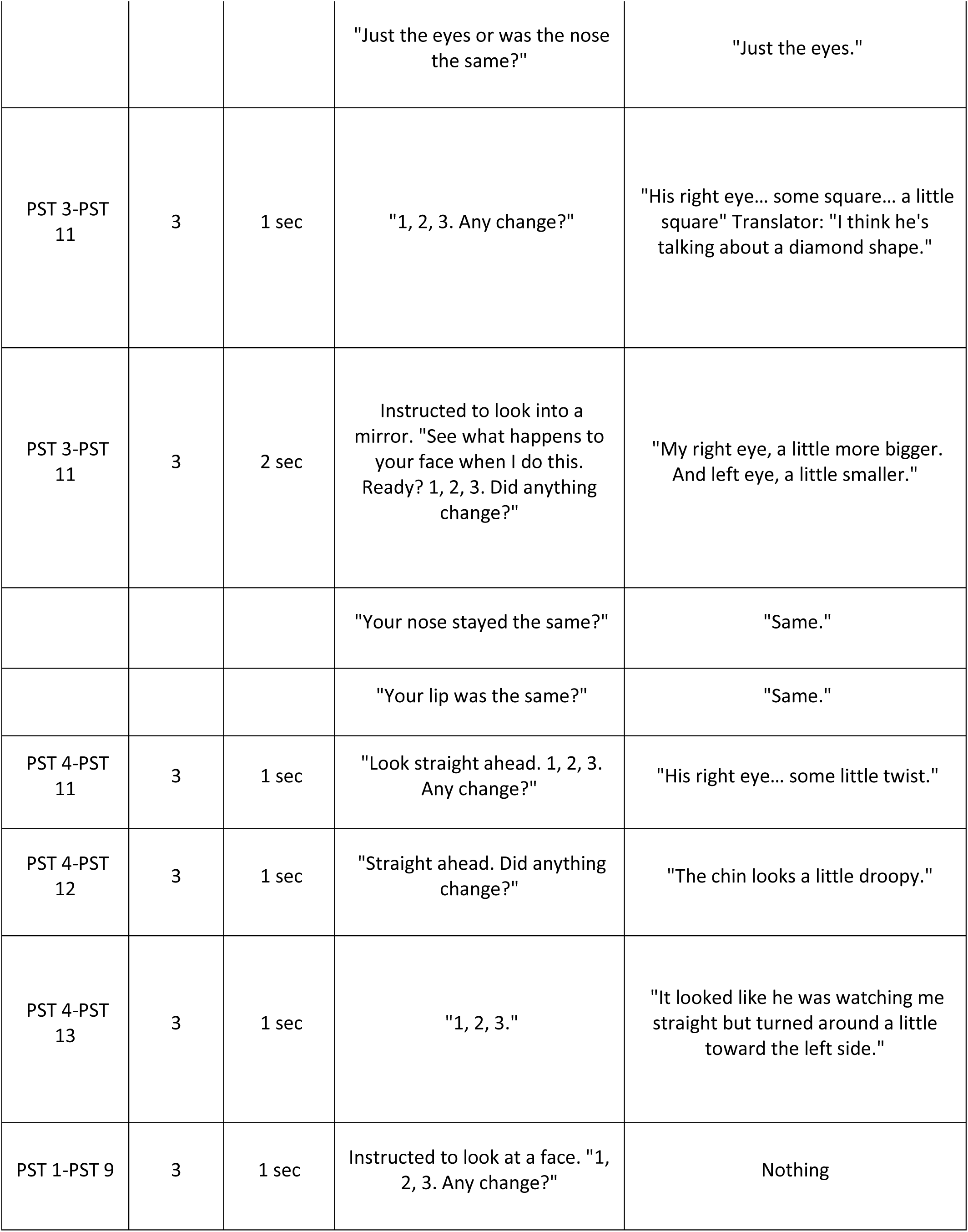
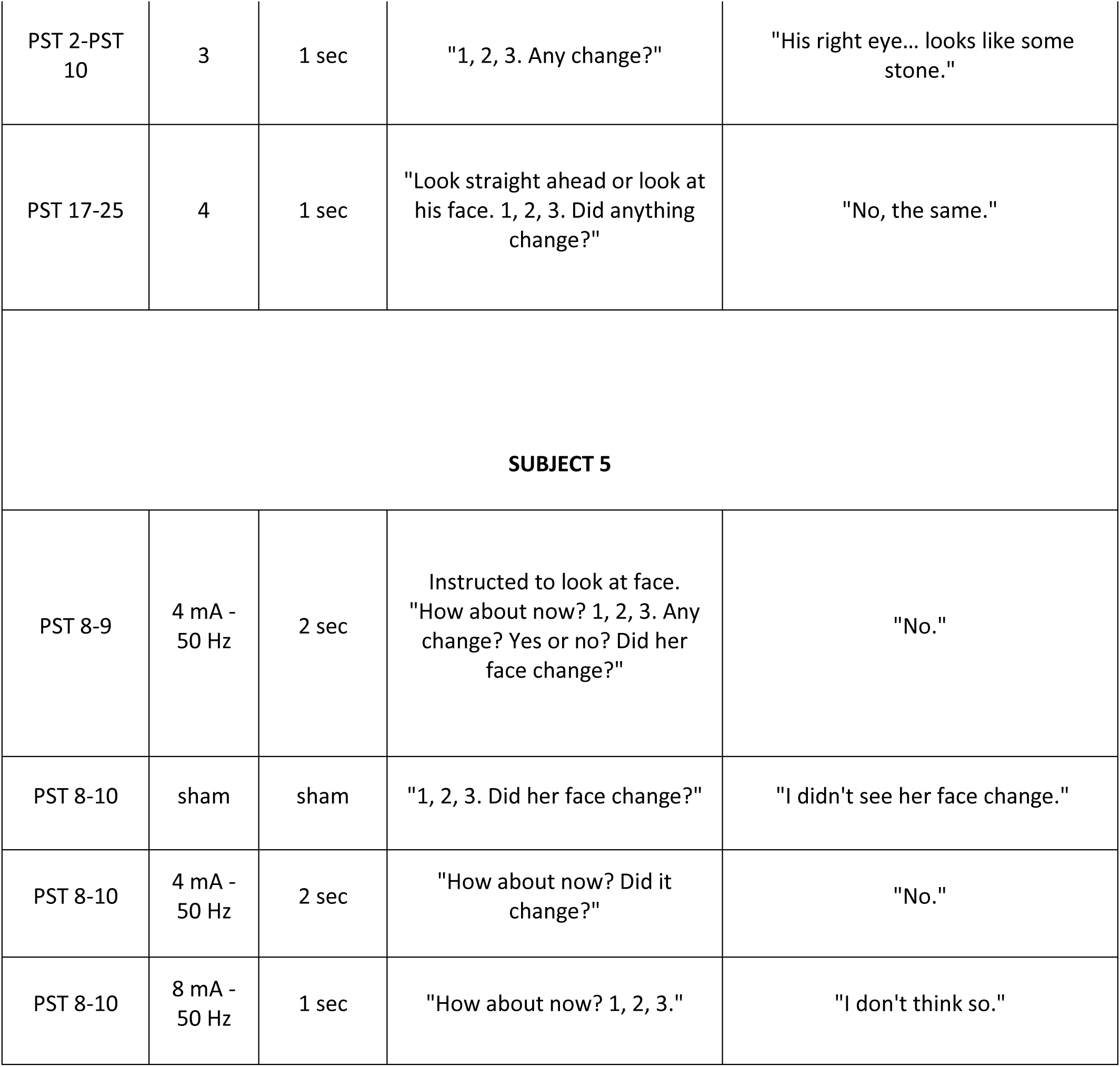
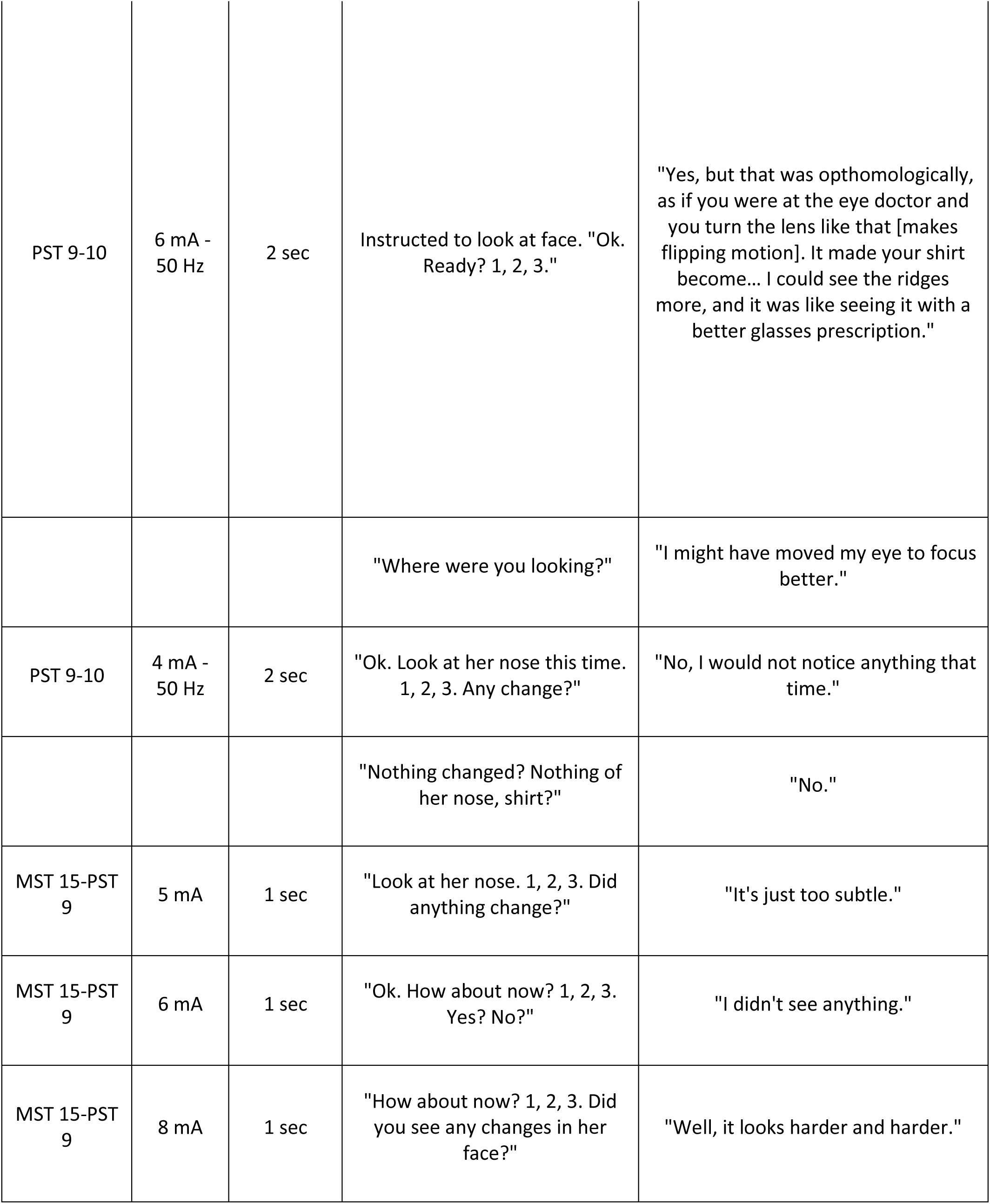
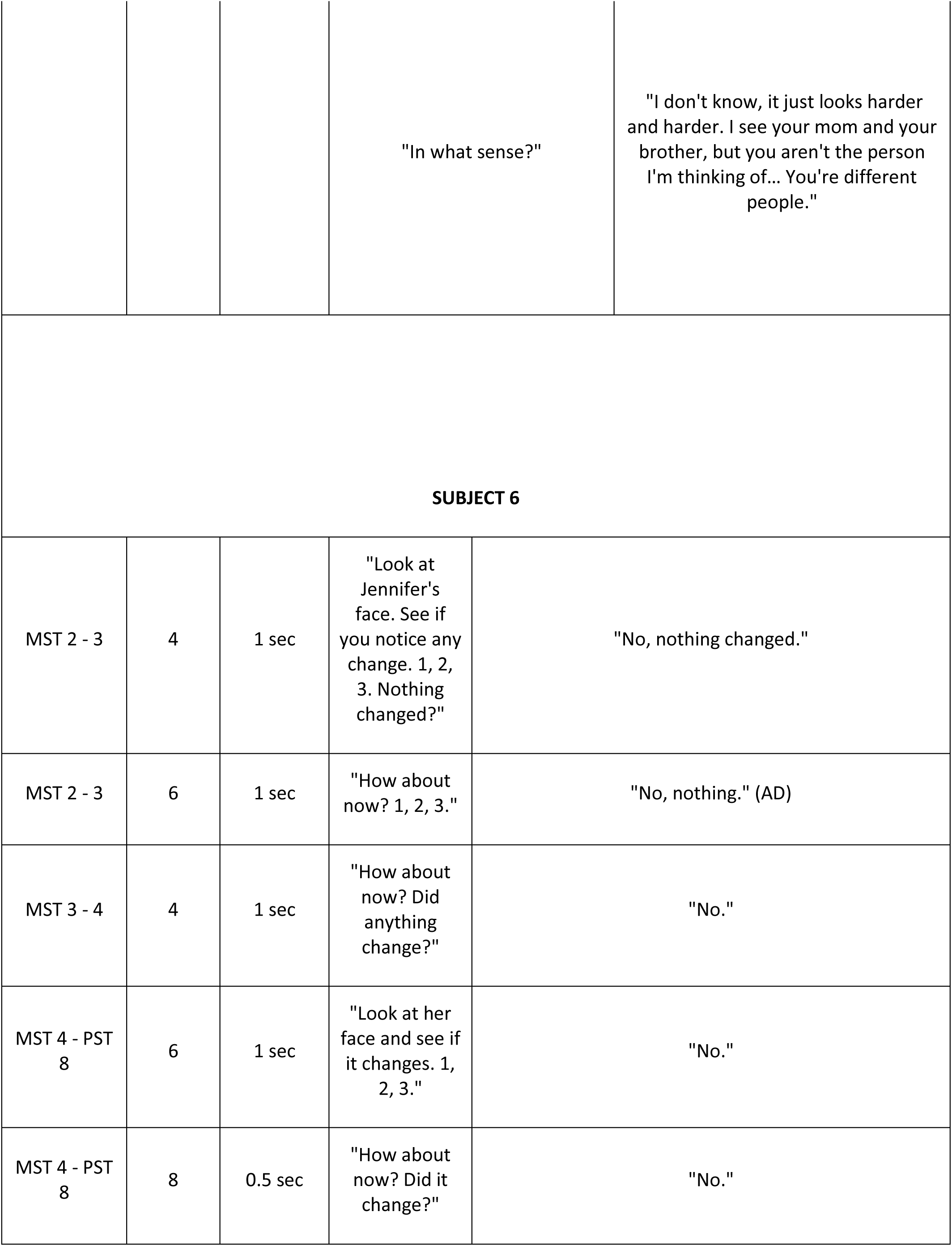
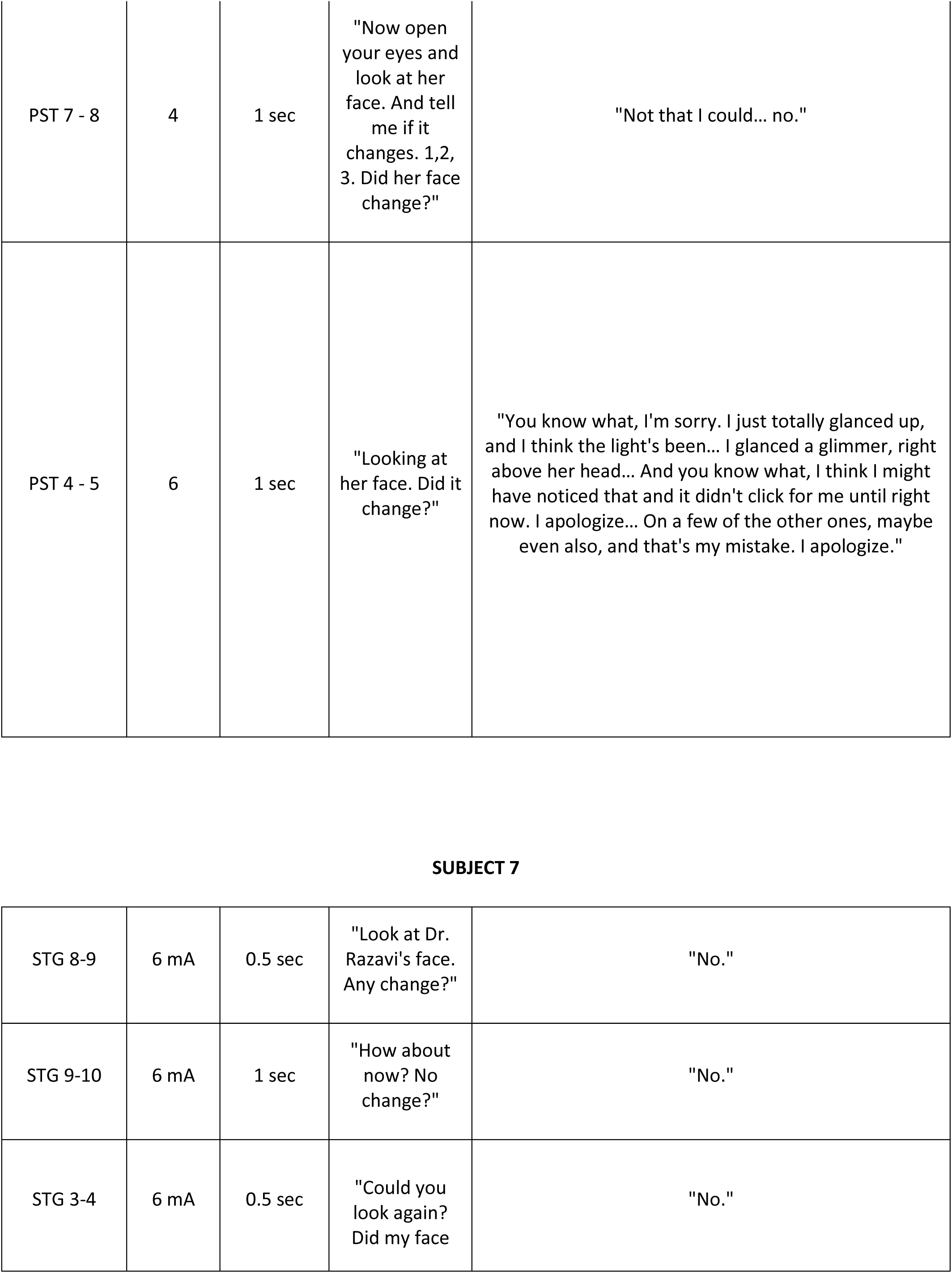
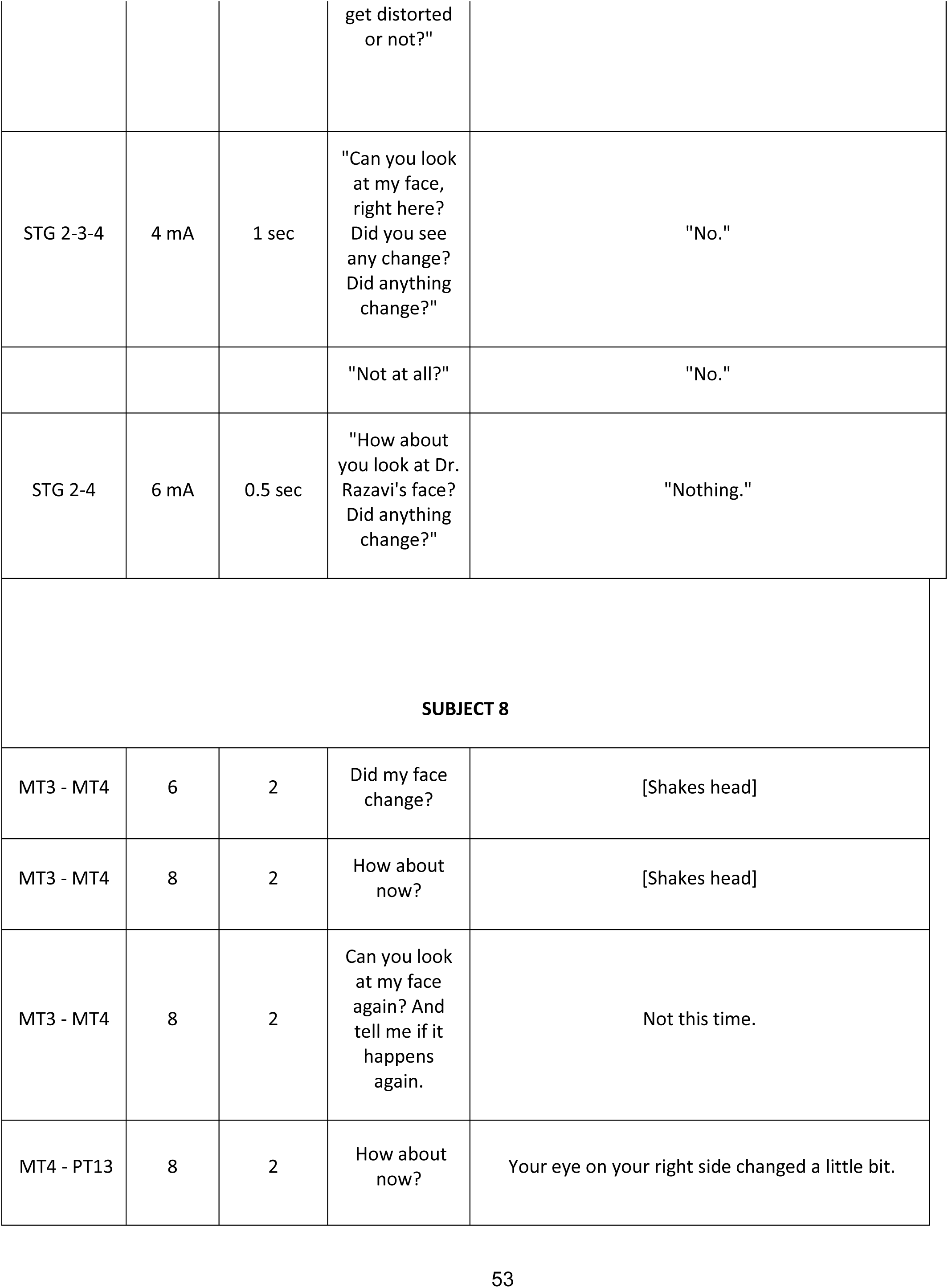
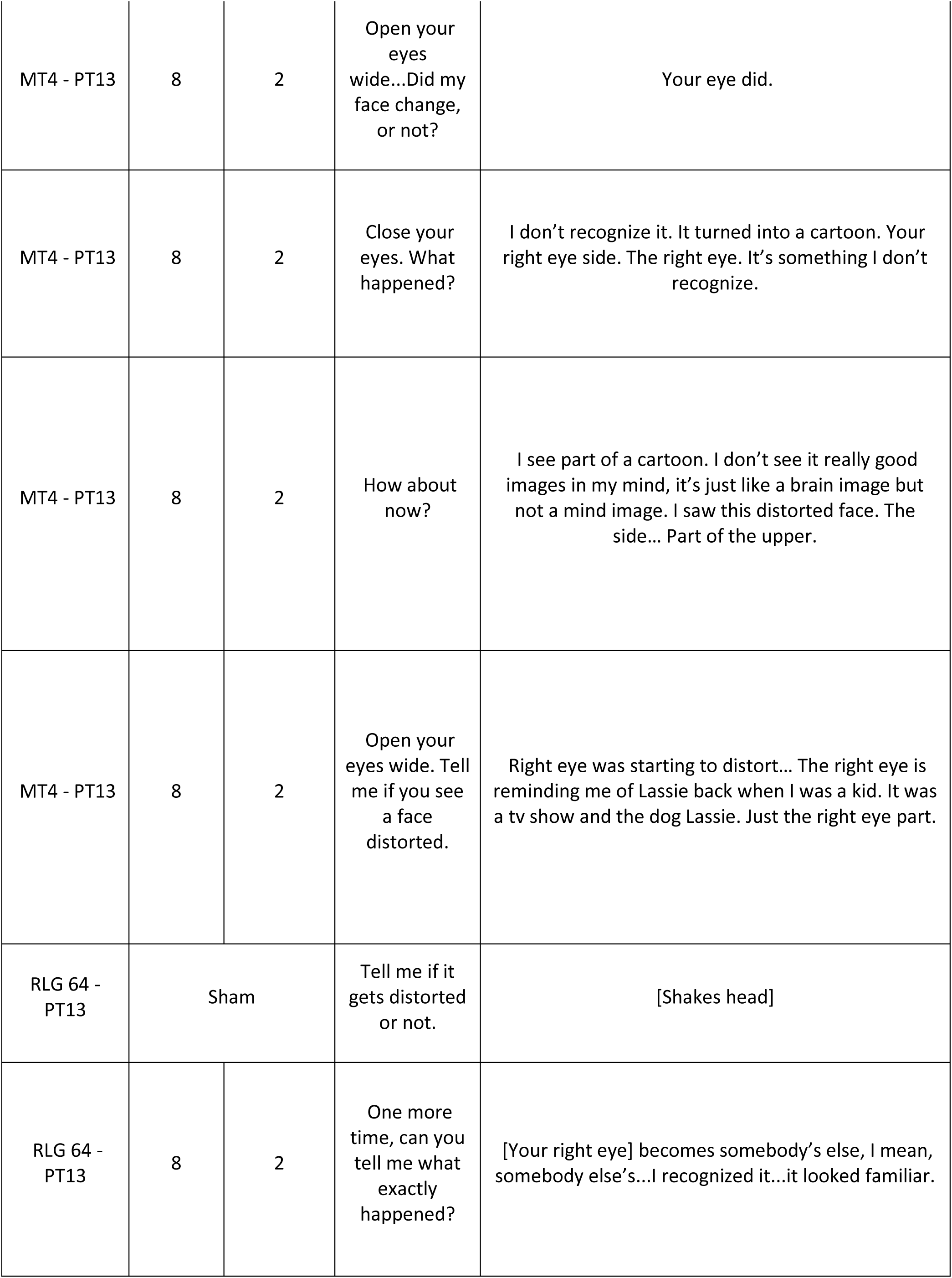
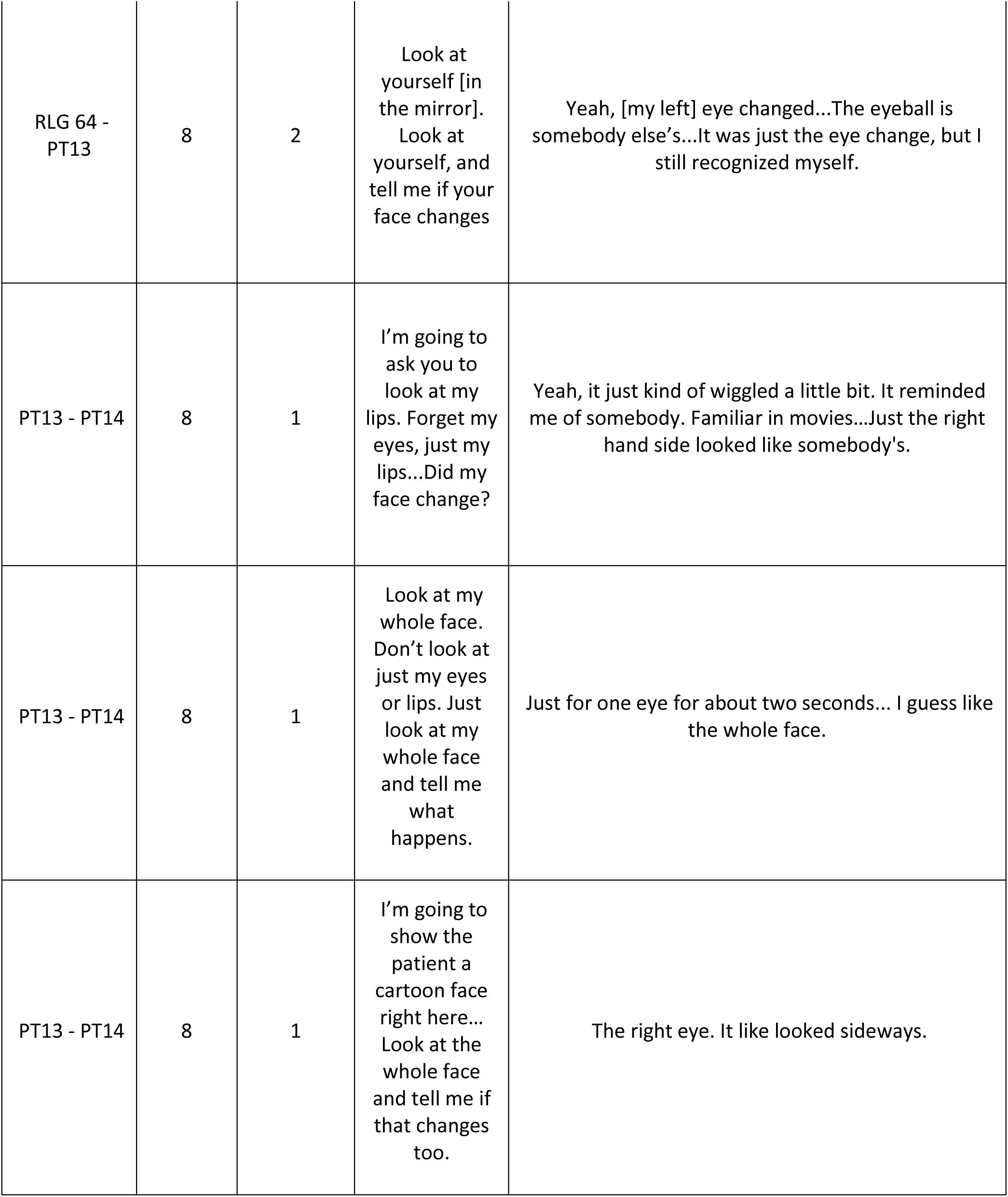
Subject responses during electrical stimulation trials.

| Subject 1 |  |  |  |  |
| --- | --- | --- | --- | --- |
| Stimulated Pair | Current (mA) | Duration (sec) | Prompt | Result |
| STG 7-STG 1 | 6 | 1.7 sec | Instructed to look at a face.<br>"How about now?" | "Nothing." |
| STG 1-STG 2 | 6 | 1.7 sec | "Changed or no?" | "No." |
| STG 3-STG 2 | 6 | 1.4 sec | Instructed to look at hand in front of face. "1, 2, 3." | "Nothing." |
| STG 3-STG 2 | 6 | 1.4 sec | "Face. 1, 2, 3." | "Nothing." |
| STG 2-STG 8 | 6 | 1.5 sec | Instructed to look at face.<br>"How about now?" | "It changed a little bit." |
|  |  |  | "Can you describe it?" | "Just that one side changed." |
| STG 4-STG 3 | 6 | 1.1 sec | "How about face?" | "Nothing." |
| STG 9-STG 3 | 6 | 1.5 sec | "Look at her face. 1, 2, 3." | "Nothing." |
| STG 10-STG 4 | 6 | 1.5 sec | "Now face. 1, 2, 3." | "No." |
| STG 7-STG 8 | 6 | 1.7 sec | "Ok, now?" | "I didn't see anything." |
| STG 13-STG 7 | 6 | 1.6 sec | "Now look at her face. Did anything change?" | "Nothing." |
| STG 9-STG 10 | 6 | 1.6 sec | "Ok. Now you are going to look at the face. Ready? 1, 2, 3." | "Nothing." |
| STG 9-STG 10 | Sham | Sham | "One more time. 1, 2, 3. Nothing?" | Nothing |
| STG 9-STG 15 | 6 | 1.6 sec | "Now look at her face. 1, 2, 3." | "No." |
| STG 10-STG 16 | 6 | 1.4 sec | "How about the face? 1, 2, 3." | "Nothing." |
| STG 14-13 | 6 | 1.5 sec | "Can you look at her face now? 1, 2, 3. Did it change?" | "Yeah. It changed a little." |
|  |  |  | "In what sense?" | "Just that left side of her face changed." |
|  |  |  | "Was it very similar to the previous feelings?" | "That one looked like it changed a little different. It looks like my wife's cousin or something." |
| STG 19-STG 13 | 6 | 1.7 sec | "Now look at the clock. 1, 2, 3." | "I didn't really see anything." |
| STG 20-STG 14 | 6 | 2.5 sec | "Ok. Alright. Now look at her face and try to describe exactly what's happening." | "It looks like my neighbor's wife or something on one side." |
|  |  |  | "How old is your neighbor's wife?" | "Uh, forty-something." |
|  |  |  | "So, you think she was looking older?" | "No. She wasn't looking older?" |
|  |  |  | "But you could still know who she was?" | "I had an idea." |
| STG 14-STG 15 | 6 | 1.8 sec | "How about now?" | "That one was like, eh, it was like a face in a movie I've seen but I can't remember the movie, just the half part [changed]." |
|  |  |  | "Let me ask you this question. Were you seeing a face or did the face change to something non-face?" | "No. It changed to a face." |
|  |  |  | "It changed to another face?" | "Yeah. Just half of it changed to another face." |
| STG 15-STG 16 | 6 | 1.6 sec | "How about now?" | "About the same thing. It looks like a person I've seen in a movie." |
|  |  |  | "Person you've seen in a movie?" | "Yeah." |
| STG 15-STG 21 | 6 | 1.7 sec | "Look straight ahead. 1, 2, 3. Did anything change?" | "Very little [changed], not too much." |
|  |  |  | "What did change?" | "Same thing. It was just that one side." |
|  |  |  | "Ok. Compared to the previous one?" | "Not as much as the previous one." |
|  |  |  | "Was the intensity different or was it the character that was different?" | "It was the same person or something that I had seen in movies before." |
|  |  |  | "Why was it less?" | "Oh, I don't know, but it was less." |
| STG 19-STG 20 | 4 | 1.5 sec | "Ok. Look at my face. How about now? Did you see any change?" | "Yeah, it changed. That side [the pt's right side] changed." |
|  |  |  | "Tell me more." | "It stayed the same size and everything. It just changed a little bit on that one side." |
|  |  |  | "Was it shaking... Did I turn into somebody else?" | "It looked like that side [PT's right side] turned into somebody else." |
|  |  |  | "Someone you have known before or what?" | "Nah. I didn't really recognize him." |
| STG 20-STG 21 | 6 | 1.7 sec | "Look at her face again one more time and let's see if it changes. You're looking at her face? And that's normal looking? How about now?" | "It changed. I couldn't pinpoint the person who it was." |
|  |  |  | "It was a familiar face?" | "It was like somebody that I knew, that I've seen, just on the left side." |
|  |  |  | "It was a familiar face? Somebody you had seen before?" | "Yeah." |
|  |  |  | "But it wasn't her face?" | "No." |
|  |  |  | "Can you say a little bit more?" | "I can't really pinpoint exactly who it was..." |
|  |  |  | "Were the eyes in the same place?" | "Yeah. Same place." |
|  |  |  | "Did the face kind of change to a different shape?" | "No. It pretty much stayed the same?" |
|  |  |  | "Male or female?" | "That time it wa male." |
|  |  |  | "But someone familiar?" | "Someone familiar." |
| **SUBJECT 3** |  |  |  |  |
| OC 26-27 | 7 | 1 sec | "Can you look at our face instead? Any change?" | "No." |
| OC 27-28 | 7 | 1 sec | "Look at my face. Any change?" | "No." |
| OC 28-29 | 7 | 1 sec | "Look at my face. No change?" | Nothing |
| **SUBJECT 4** |  |  |  |  |
| PST 1-PST 9 | 3 | 1 sec | Instructed to look at a face. "1, 2, 3. Any change?" | Nothing |
| PST 2-PST 3<br>& PST 11-PST 12 | 4 | 1 sec | "Ok, how about now? 1, 2, and 3." | "The same... the right eyes cross... it's not a circle... it" |
|  |  |  | "Can you draw what happened?" | *draws what happened, looks like a square* |
|  |  |  | "But the whole face is the same?" | "The whole face, not much changed." |
| PST 2-PST 3<br>& PST 11-PST 12 | 4 | 1 sec | "Concentrate on the whole face. Ready? One last time. 1, 2, 3. Did anything change?" | "Just eyes." |
|  |  |  | "Just eyes?" | "Same as before." |
| PST 2-PST<br>10 | 3 | 1 sec | "1, 2, 3. Any change?" | "His right eye... looks like some stone." |
| PST 2-PST<br>11 | 4 | 2 sec | "Look at her face and tell me what you notice. Just look straight ahead and tell me what happens. Did anything change? Explain." | "Her right eye looks like a dance motion." |
|  |  |  | "Was the whole face the same?" | "I didn't see the whole face."<br>Translator: "Initially he was concentrating on her right eye but the face also had some fading." |
|  |  |  | "Was the face distorted?" | "The whole face stays about the same... only the eyes change." |
| PST 2-PST 12 | 4 | 1 sec | "Look straight ahead. Look at his nose and pay attention to his face. 1, 2, 3." | "No change, but there was some twist [gesturing to bridge of nose]." |
| PST 2-PST 3 & PST 11-PST 12 | Sham | Sham | "Just look at her. Ready? 1, 2, and 3. Anything change?" | "No, it's not changed." |
| PST 3-PST 11 | 4 | 1 sec | "Just look at his face. Ok, go." | "[The pt] was looking at my eye and he said that initially it looked like a circle but then it turned into a rectangular shape." |
|  |  |  | "Did the whole face remain the same or did it change?" | [No answer] |
| PST 3-PST 11 | 4 | 2 sec | "How about now? Look straight ahead to his face." | "The right eye looked normal and then it twisted." |
|  |  |  | "Just the eyes or was the nose the same?" | "Just the eyes." |
| PST 3-PST 11 | 3 | 1 sec | "1, 2, 3. Any change?" | "His right eye... some square... a little square" Translator: "I think he's talking about a diamond shape." |
| PST 3-PST 11 | 3 | 2 sec | Instructed to look into a mirror. "See what happens to your face when I do this. Ready? 1, 2, 3. Did anything change?" | "My right eye, a little more bigger. And left eye, a little smaller." |
|  |  |  | "Your nose stayed the same?" | "Same." |
|  |  |  | "Your lip was the same?" | "Same." |
| PST 4-PST 11 | 3 | 1 sec | "Look straight ahead. 1, 2, 3. Any change?" | "His right eye... some little twist." |
| PST 4-PST 12 | 3 | 1 sec | "Straight ahead. Did anything change?" | "The chin looks a little droopy." |
| PST 4-PST 13 | 3 | 1 sec | "1, 2, 3." | "It looked like he was watching me straight but turned around a little toward the left side." |
| PST 1-PST 9 | 3 | 1 sec | Instructed to look at a face. "1, 2, 3. Any change?" | Nothing |
| PST 2-PST<br>10 | 3 | 1 sec | "1, 2, 3. Any change?" | "His right eye... looks like some<br>stone." |
| PST 17-25 | 4 | 1 sec | "Look straight ahead or look at<br>his face. 1, 2, 3. Did anything<br>change?" | "No, the same." |
| **SUBJECT 5** |  |  |  |  |
| PST 8-9 | 4 mA -<br>50 Hz | 2 sec | Instructed to look at face.<br>"How about now? 1, 2, 3. Any<br>change? Yes or no? Did her<br>face change?" | "No." |
| PST 8-10 | sham | sham | "1, 2, 3. Did her face change?" | "I didn't see her face change." |
| PST 8-10 | 4 mA -<br>50 Hz | 2 sec | "How about now? Did it<br>change?" | "No." |
| PST 8-10 | 8 mA -<br>50 Hz | 1 sec | "How about now? 1, 2, 3." | "I don't think so." |
| PST 9-10 | 6 mA -<br>50 Hz | 2 sec | Instructed to look at face. "Ok.<br>Ready? 1, 2, 3." | "Yes, but that was opthomologically,<br>as if you were at the eye doctor and<br>you turn the lens like that [makes<br>flipping motion]. It made your shirt<br>become... I could see the ridges<br>more, and it was like seeing it with a<br>better glasses prescription." |
|  |  |  | "Where were you looking?" | "I might have moved my eye to focus<br>better." |
| PST 9-10 | 4 mA -<br>50 Hz | 2 sec | "Ok. Look at her nose this time.<br>1, 2, 3. Any change?" | "No, I would not notice anything that<br>time." |
|  |  |  | "Nothing changed? Nothing of<br>her nose, shirt?" | "No." |
| MST 15-PST<br>9 | 5 mA | 1 sec | "Look at her nose. 1, 2, 3. Did<br>anything change?" | "It's just too subtle." |
| MST 15-PST<br>9 | 6 mA | 1 sec | "Ok. How about now? 1, 2, 3.<br>Yes? No?" | "I didn't see anything." |
| MST 15-PST<br>9 | 8 mA | 1 sec | "How about now? 1, 2, 3. Did<br>you see any changes in her<br>face?" | "Well, it looks harder and harder." |
|  |  |  | "In what sense?" | "I don't know, it just looks harder and harder. I see your mom and your brother, but you aren't the person I'm thinking of... You're different people." |
| **SUBJECT 6** |  |  |  |  |
| MST 2 - 3 | 4 | 1 sec | "Look at Jennifer's face. See if you notice any change. 1, 2, 3. Nothing changed?" | "No, nothing changed." |
| MST 2 - 3 | 6 | 1 sec | "How about now? 1, 2, 3." | "No, nothing." (AD) |
| MST 3 - 4 | 4 | 1 sec | "How about now? Did anything change?" | "No." |
| MST 4 - PST<br>8 | 6 | 1 sec | "Look at her face and see if it changes. 1, 2, 3." | "No." |
| MST 4 - PST<br>8 | 8 | 0.5 sec | "How about now? Did it change?" | "No." |
| PST 7 - 8 | 4 | 1 sec | "Now open your eyes and look at her face. And tell me if it changes. 1,2, 3. Did her face change?" | "Not that I could... no." |
| PST 4 - 5 | 6 | 1 sec | "Looking at her face. Did it change?" | "You know what, I'm sorry. I just totally glanced up, and I think the light's been... I glanced a glimmer, right above her head... And you know what, I think I might have noticed that and it didn't click for me until right now. I apologize... On a few of the other ones, maybe even also, and that's my mistake. I apologize." |
| SUBJECT 7 |  |  |  |  |
| STG 8-9 | 6 mA | 0.5 sec | "Look at Dr. Razavi's face. Any change?" | "No." |
| STG 9-10 | 6 mA | 1 sec | "How about now? No change?" | "No." |
| STG 3-4 | 6 mA | 0.5 sec | "Could you look again? Did my face get distorted or not?" | "No." |
| STG 2-3-4 | 4 mA | 1 sec | "Can you look at my face, right here? Did you see any change? Did anything change?" | "No." |
|  |  |  | "Not at all?" | "No." |
| STG 2-4 | 6 mA | 0.5 sec | "How about you look at Dr. Razavi's face? Did anything change?" | "Nothing." |
| **SUBJECT 8** |  |  |  |  |
| MT3 - MT4 | 6 | 2 | Did my face change? | [Shakes head] |
| MT3 - MT4 | 8 | 2 | How about now? | [Shakes head] |
| MT3 - MT4 | 8 | 2 | Can you look at my face again? And tell me if it happens again. | Not this time. |
| MT4 - PT13 | 8 | 2 | How about now? | Your eye on your right side changed a little bit. |
| MT4 - PT13 | 8 | 2 | Open your eyes wide...Did my face change, or not? | Your eye did. |
| MT4 - PT13 | 8 | 2 | Close your eyes. What happened? | I don't recognize it. It turned into a cartoon. Your right eye side. The right eye. It's something I don't recognize. |
| MT4 - PT13 | 8 | 2 | How about now? | I see part of a cartoon. I don't see it really good images in my mind, it's just like a brain image but not a mind image. I saw this distorted face. The side... Part of the upper. |
| MT4 - PT13 | 8 | 2 | Open your eyes wide. Tell me if you see a face distorted. | Right eye was starting to distort... The right eye is reminding me of Lassie back when I was a kid. It was a tv show and the dog Lassie. Just the right eye part. |
| RLG 64 - PT13 | Sham |  | Tell me if it gets distorted or not. | [Shakes head] |
| RLG 64 - PT13 | 8 | 2 | One more time, can you tell me what exactly happened? | [Your right eye] becomes somebody's else, I mean, somebody else's...I recognized it...it looked familiar. |
| RLG 64 - PT13 | 8 | 2 | Look at yourself [in the mirror].<br>Look at yourself, and tell me if your face changes | Yeah, [my left] eye changed...The eyeball is somebody else's...It was just the eye change, but I still recognized myself. |
| PT13 - PT14 | 8 | 1 | I'm going to ask you to look at my lips. Forget my eyes, just my lips...Did my face change? | Yeah, it just kind of wiggled a little bit. It reminded me of somebody. Familiar in movies...Just the right hand side looked like somebody's. |
| PT13 - PT14 | 8 | 1 | Look at my whole face. Don't look at just my eyes or lips. Just look at my whole face and tell me what happens. | Just for one eye for about two seconds... I guess like the whole face. |
| PT13 - PT14 | 8 | 1 | I'm going to show the patient a cartoon face right here... Look at the whole face and tell me if that changes too. | The right eye. It like looked sideways. |

**Author contributions**
J.P. and V.R. contributed to the design of experiments and data acquisition; J.S., O.R., J.M.M. and A.L.D developed analysis tools and all authors contributed to data analysis and writing of the manuscript

## REFERENCES

1. Damasio AR, Damasio H, Van Hoesen GW. Prosopagnosia: anatomic basis and behavioral mechanisms. Neurology. 1982;32(4):331–41.

2. Meadows JC. The anatomical basis of prosopagnosia. Journal of neurology, neurosurgery, and psychiatry. 1974;37(5):489–501.

3. Sergent J, Ohta S, MacDonald B. Functional neuroanatomy of face and object processing. A positron emission tomography study. Brain : a journal of neurology. 1992;115 Pt 1:15–36.

4. Puce A, Allison T, Gore JC, McCarthy G. Face-sensitive regions in human extrastriate cortex studied by functional MRI. J Neurophysiol. 1995;74(3):1192–9.

5. Kanwisher N, McDermott J, Chun MM. The fusiform face area: a module in human extrastriate cortex specialized for face perception. The Journal of neuroscience : the official journal of the Society for Neuroscience. 1997;17(11):4302–11.

6. Haxby JV, Ungerleider LG, Horwitz B, Maisog JM, Rapoport SI, Grady CL. Face encoding and recognition in the human brain. Proc Natl Acad Sci U S A. 1996;93(2):922–7.

7. Bentin S, Allison T, Puce A, Perez E, McCarthy G. Electrophysiological Studies of Face Perception in Humans. J Cogn Neurosci. 1996;8(6):551–65.

8. Vida MD, Nestor A, Plaut DC, Behrmann M. Spatiotemporal dynamics of similarity-based neural representations of facial identity. Proc Natl Acad Sci U S A. 2017;114(2):388–93.

9. Yang Y, Xu Y, Jew CA, Pyles JA, Kass RE, Tarr MJ. Exploring the spatio-temporal neural basis of face learning. J Vis. 2017;17(6):1.

10. Bruce C, Desimone R, Gross CG. Visual properties of neurons in a polysensory area in superior temporal sulcus of the macaque. J Neurophysiol. 1981;46(2):369–84.

11. Kendrick KM, Baldwin BA. Cells in temporal cortex of conscious sheep can respond preferentially to the sight of faces. Science. 1987;236(4800):448–50.

12. Ungerleider LG, Haxby JV. ‘What’ and ‘where’ in the human brain. Curr Opin Neurobiol. 1994;4(2):157–65.

13. Tsao DY, Freiwald WA, Knutsen TA, Mandeville JB, Tootell RB. Faces and objects in macaque cerebral cortex. Nat Neurosci. 2003;6(9):989–95.

14. Afraz SR, Kiani R, Esteky H. Microstimulation of inferotemporal cortex influences face categorization. Nature. 2006;442(7103):692–5.

15. DiCarlo JJ, Zoccolan D, Rust NC. How does the brain solve visual object recognition? Neuron. 2012;73(3):415–34.

16. Behrmann M, Plaut DC. Distributed circuits, not circumscribed centers, mediate visual recognition. Trends Cogn Sci. 2013;17(5):210–9.

17. Haxby JV, Gobbini MI, Furey ML, Ishai A, Schouten JL, Pietrini P. Distributed and overlapping representations of faces and objects in ventral temporal cortex. Science. 2001;293(11577229):2425–30.

18. Parvizi J, Kastner DB. Human intracranial EEG: Promises and Limitations. Nat Neurosci. 2018(21):474–83

19. Jonas J, Rossion B, Brissart H, Frismand S, Jacques C, Hossu G, et al. Beyond the core face-processing network: Intracerebral stimulation of a face-selective area in the right anterior fusiform gyrus elicits transient prosopagnosia. Cortex. 2015;72:140–55.

20. Ghuman AS, Brunet NM, Li Y, Konecky RO, Pyles JA, Walls SA, et al. Dynamic encoding of face information in the human fusiform gyrus. Nat Commun. 2014;5:5672.

21. Davidesco I, Zion-Golumbic E, Bickel S, Harel M, Groppe DM, Keller CJ, et al. Exemplar selectivity reflects perceptual similarities in the human fusiform cortex. Cereb Cortex. 2014;24(7):1879–93.

22. Keller CJ, Davidesco I, Megevand P, Lado FA, Malach R, Mehta AD. Tuning face perception with electrical stimulation of the fusiform gyrus. Hum Brain Mapp. 2017.

23. Hamame CM, Vidal JR, Perrone-Bertolotti M, Ossandon T, Jerbi K, Kahane P, et al. Functional selectivity in the human occipitotemporal cortex during natural vision: evidence from combined intracranial EEG and eye-tracking. Neuroimage. 2014;95:276–86.

24. Murphey DK, Maunsell JH, Beauchamp MS, Yoshor D. Perceiving electrical stimulation of identified human visual areas. Proc Natl Acad Sci U S A. 2009;106(13):5389–93.

25. Jacques C, Witthoft N, Weiner KS, Foster BL, Rangarajan V, Hermes D, et al. Corresponding ECoG and fMRI category-selective signals in human ventral temporal cortex. Neuropsychologia. 2016;83:14–28.

26. Rangarajan V, Parvizi J. Functional asymmetry between the left and right human fusiform gyrus explored through electrical brain stimulation. Neuropsychologia. 2016;83:29–36.

27. Rangarajan V, Hermes D, Foster BL, Weiner KS, Jacques C, Grill-Spector K, et al. Electrical stimulation of the left and right human fusiform gyrus causes different effects in conscious face perception. J Neurosci. 2014;34(38):12828–36.

28. Parvizi J, Jacques C, Foster BL, Withoft N, Rangarajan V, Weiner KS, et al. Electrical stimulation of human fusiform face-selective regions distorts face perception. The Journal of neuroscience : the official journal of the Society for Neuroscience. 2012;32(43):14915–20.

29. Rakotomamonjy A, Bach F, Canu S, Grandvalet Y. Simple MKL. Journal of Machine Learning. 2008(9):2491–521.

30. Schrouff J, Mourao-Miranda J, Phillips C, Parvizi J. Decoding intracranial EEG data with multiple kernel learning method. J Neurosci Methods. 2016;261:19–28.

31. Canolty RT, Knight RT. The functional role of cross-frequency coupling. Trends in Cognitive Sciences. 2010;14(11):506–15.

32. Crone NE, Miglioretti DL, Gordon B, Lesser RP. Functional mapping of human sensorimotor cortex with electrocorticographic spectral analysis. II. Event-related synchronization in the gamma band. Brain. 1998;121(Pt 12):2301–15.

33. Nir Y, Fisch L, Mukamel R, Gelbard-Sagiv H, Arieli A, Fried I, et al. Coupling between neuronal firing rate, gamma LFP, and BOLD fMRI is related to interneuronal correlations. Curr Biol. 2007;17(17686438):1275–85.

34. Nir Y, Mukamel R, Dinstein I, Privman E, Harel M, Fisch L, et al. Interhemispheric correlations of slow spontaneous neuronal fluctuations revealed in human sensory cortex. Nat Neurosci. 2008;11(9):1100–8.

35. Mukamel R. Coupling Between Neuronal Firing, Field Potentials, and fMRI in Human Auditory Cortex. Science. 2005;309(5736):951–4.

36. Niessing J, Ebisch B, Schmidt KE, Niessing M, Singer W, Galuske RA. Hemodynamic signals correlate tightly with synchronized gamma oscillations. Science. 2005;309(5736):948– 51.

37. Manning JR, Jacobs J, Fried I, Kahana MJ. Broadband shifts in local field potential power spectra are correlated with single-neuron spiking in humans. J Neurosci. 2009;29(43):13613–20.

38. Ray S, Crone NE, Niebur E, Franaszczuk PJ, Hsiao SS. Neural Correlates of High-Gamma Oscillations (60-200 Hz) in Macaque Local Field Potentials and Their Potential Implications in Electrocorticography. J Neurosci. 2008;28(45):11526–36.

39. Logothetis NK, Pauls J, Augath M, Trinath T, Oeltermann A. Neurophysiological investigation of the basis of the fMRI signal. Nature. 2001;412(11449264):150–7.

40. Goense JB, Logothetis NK. Neurophysiology of the BOLD fMRI signal in awake monkeys. Curr Biol. 2008;18(9):631–40. Epub 2008 Apr 24.

41. Kreiman G, Hung CP, Kraskov A, Quiroga RQ, Poggio T, DiCarlo JJ. Object selectivity of local field potentials and spikes in the macaque inferior temporal cortex. Neuron. 2006;49(3):433–45.

42. Liu J, Newsome WT. Local field potential in cortical area MT: stimulus tuning and behavioral correlations. J Neurosci. 2006;26(30):7779–90.

43. Ray S, Maunsell JH. Different origins of gamma rhythm and high-gamma activity in macaque visual cortex. PLoS Biol. 2011;9(4):e1000610.

44. Park SH, Russ BE, McMahon DBT, Koyano KW, Berman RA, Leopold DA. Functional Subpopulations of Neurons in a Macaque Face Patch Revealed by Single-Unit fMRI Mapping. Neuron. 2017;95(4):971–81 e5.

45. Weichwald S, Meyer T, Ozdenizci O, Scholkopf B, Ball T, Grosse-Wentrup M. Causal interpretation rules for encoding and decoding models in neuroimaging. Neuroimage. 2015;110:48–59.

46. Shehzad Z, McCarthy G. Category representations in the brain are both discretely localized and widely distributed. J Neurophysiol. 2018;119(6):2256–64.

47. Schrouff J, Monteiro JM, Portugal L, Rosa MJ, Phillips C, Mourao-Miranda J. Embedding Anatomical or Functional Knowledge in Whole-Brain Multiple Kernel Learning Models. Neuroinformatics. 2018;16(1):117–43.

48. Jonas J, Maillard L, Frismand S, Colnat-Coulbois S, Vespignani H, Rossion B, et al. Self-face hallucination evoked by electrical stimulation of the human brain. Neurology. 2014;83(4):336–8.

49. Schalk G, Kapeller C, Guger C, Ogawa H, Hiroshima S, Lafer-Sousa R, et al. Facephenes and rainbows: Causal evidence for functional and anatomical specificity of face and color processing in the human brain. Proc Natl Acad Sci U S A. 2017;114(46):12285–90.

50. Allison T, Ginter H, McCarthy G, Nobre AC, Puce A, Luby M, et al. Face recognition in human extrastriate cortex. J Neurophysiol. 1994;71(2):821–5.

51. Puce A, Allison T, McCarthy G. Electrophysiological studies of human face perception. III: Effects of top-down processing on face-specific potentials. Cereb Cortex. 1999;9(5):445–58.

52. Moeller S, Crapse T, Chang L, Tsao DY. The effect of face patch microstimulation on perception of faces and objects. Nat Neurosci. 2017;20(5):743–52.

53. Weiner KS, Natu VS, Grill-Spector K. On object selectivity and the anatomy of the human fusiform gyrus. Neuroimage. 2018.

54. Weiner KS, Barnett MA, Lorenz S, Caspers J, Stigliani A, Amunts K, et al. The Cytoarchitecture of Domain-specific Regions in Human High-level Visual Cortex. Cereb Cortex. 2017;27(1):146–61.

55. Mesulam MM. Large-scale neurocognitive networks and distributed processing for attention, language, and memory. Ann Neurol. 1990;28(5):597–613.

56. Schrouff J, Rosa MJ, Rondina JM, Marquand AF, Chu C, Ashburner J, et al. PRoNTo: Pattern Recognition for Neuroimaging Toolbox. Neuroinformatics. 2013;11(3):319–37.

57. Hermes D, Miller KJ, Noordmans HJ, Vansteensel MJ, Ramsey NF. Automated electrocorticographic electrode localization on individually rendered brain surfaces. J Neurosci Methods. 2010;185(2):293–8.

58. Pereira F, Mitchell TM, Botvinick M. Machine learning classifiers and fMRI: a tutorial overview. Neuroimage. 2009;45:S199–S209.

59. Haufe S, Meinecke F, Gorgen K, Dahne S, Haynes JD, Blankertz B, et al. On the interpretation of weight vectors of linear models in multivariate neuroimaging. Neuroimage. 2014;87:96–110.

60. Tang H, Buia C, Madhavan R, Crone NE, Madsen JR, Anderson WS, et al. Spatiotemporal dynamics underlying object completion in human ventral visual cortex. Neuron. 2014;83(3):736–48.

61. Schrouff J, Mourao-Miranda J. Interpreting weight maps in terms of cognitive or clinical neuroscience: nonsense? International Workshop on Pattern Recognition in Neuroimaging (PRNI). Singapore 2018. p. 1–4.

62. Winawer J, Parvizi J. Linking Electrical Stimulation of Human Primary Visual Cortex, Size of Affected Cortical Area, Neuronal Responses, and Subjective Experience. Neuron. 2016;92(6):1213–9.

